# LIN-23 Affects *C. elegans* Pathogen and Stress Resistance by Modulating SKN-1 Activity

**DOI:** 10.64898/2026.07.22.740106

**Authors:** Larissa A. Tavizón, Carolaing Gabaldón, Melissa R. Cruz, Olivia Munsey, Danielle A. Garsin

## Abstract

During pathogen infection, the *C. elegans* transcription factor SKN-1 is activated through the p38 MAPK cascade to protect against oxidative damage and promote host survival. SKN-1, the functional ortholog of the mammalian Nrf family of transcription factors, participates in various biological processes and is subject to complex regulation. In this study, we identify a previously unrecognized role for LIN-23 in regulating SKN-1 during adult stress conditions. LIN-23 is an F-box protein that functions as the substrate recognition component of the Skp–Cullin–F-box (SCF) E3 ubiquitin ligase complex and has been implicated in diverse cellular processes, including cell cycle regulation, neurite outgrowth, and centrosome duplication. Although LIN-23 has previously been reported to negatively regulate SKN-1 in other contexts, our findings demonstrate that during pathogen exposure, LIN-23 acts as a positive regulator of SKN-1 activity. Specifically, loss of LIN-23 reduced SKN-1 activity and decreased survival of infected adult animals. Having established this novel relationship between LIN-23 and SKN-1, we investigated the mechanism by which LIN-23 regulates SKN-1 activity. Because SKN-1 activation occurs via the p38 MAPK signaling pathway, followed by nuclear localization, we examined whether LIN-23 influences these events, but observed no decrease in p38 MAPK phosphorylation or SKN-1 nuclear localization. Instead, we provide evidence that LIN-23 function is dependent on WDR-23, a well-established negative regulator of SKN-1. A model is proposed in which LIN-23 promotes SKN-1 activity by targeting nuclear WDR-23 for degradation.

**SUMMARY:** SKN-1 is a *C. elegans* transcription factor and the ortholog of mammalian Nrf proteins. Under oxidative stress conditions, including those induced by infection, SKN-1 plays a protective role. Since SKN-1 regulation is complex, understanding the mechanisms that modulate its activity is important for defining stress response pathways. The authors demonstrate that the F-box protein LIN-23 functions as a positive regulator of SKN-1. Their genetic analyses indicate that LIN-23 does not influence the canonical SKN-1 activation cascade. Instead, LIN-23 appears to regulate SKN-1 activity by modulating a negative regulator, WDR-23.

## INTRODUCTION

During infection, one of the most significant physiological responses of the innate immune system is inflammation. This host defense mechanism activates signaling pathways essential for combating harmful pathogens; however, persistent inflammation has been linked to tissue damage and disease (Bender et al., 2025; Yacine et al., 2025). These detrimental effects are attributable to an imbalance between the production and clearance of reactive oxygen species (ROS) that occurs during inflammatory responses, resulting in oxidative stress (Mittal et al., 2014; Pizzino et al., 2017). In mammals, the Nrf family of transcription factors mediate responses to stress to maintain cellular homeostasis, with Nrf2 serving as a central regulator of antioxidant defense (He et al., 2020; Saha et al., 2020; Ngo & Duennwald, 2022). Similarly, when infected, the model organism *Caenorhabditis elegans* generates ROS through the Ce-Duox-1/BLI-3 dual oxidase, while a detoxification response, regulated by the transcription factor SKN-1, the functional ortholog of mammalian Nrf2 (Hoeven et al., 2011), limits collateral damage.

SKN-1 is central to multiple protective responses in *C. elegans* and is activated by diverse stimuli (Li et al., 2011; Glover-Cutter et al., 2013; Steinbaugh et al., 2015; Blackwell et al., 2015). It plays a critical role in regulating stress-responsive gene expression under a variety of conditions, including pathogen infection. During infection with intestinal pathogens such as *Enterococcus faecalis* and *Pseudomonas aeruginosa*, SKN-1 is activated through the p38 MAPK signaling cascade, leading to induction of the detoxification response (Hoeven et al., 2011; Papp et al., 2012). Consistent with this role, previous studies have demonstrated reduced survival in *skn-1* mutants following pathogen exposure, indicating that SKN-1 is required for effective host defense (Hoeven et al., 2011; Papp et al., 2012). The regulatory network controlling SKN-1-mediated protection against infection is complex and includes diverse factors, such as the tribbles pseudokinase NIPI-3 and the AAA ATPase CDC-48, which affect SKN-1 activity by modulating the p38 MAPK signaling pathway (Gabaldón et al., 2024; C. Wu et al., 2021).

Another important regulator of SKN-1 is WDR-23, a WD-40 repeat-containing protein that serves as a scaffold for protein-protein interactions in the CUL4/DDB1 ubiquitin ligase. WDR-23 was previously established as a negative regulator of SKN-1 in the context of stress induced by chemical oxidants (Choe et al., 2009; Tang & Choe, 2015) and pathogen exposure (Hoeven et al., 2011; Papp et al., 2012). More recent work showed that different isoforms differentially affect SKN-1 regulation. WDR-23A is reported to positively regulate SKN-1, whereas WDR-23B is reported to negatively regulate SKN-1 (Spatola et al., 2019). Furthermore, these isoforms have distinct cellular localizations: WDR-23A is cytoplasmic, and WDR-23B is nuclear (Lo et al., 2017; Staab et al., 2013). Finally, the WDR-23B isoform complemented the oxidative stress phenotype associated with WDR-23 loss, whereas WDR-23A did not (Spatola et al., 2019).

In this work, we report the identification of another factor that modulates SKN-1 activity during infection: the F-box protein LIN-23. Previous work on LIN-23 characterized it as a substrate recognition subunit of the Skp–Cullin–F-box (SCF) complex, binding specific target proteins for ubiquitination (Kipreos et al., 2000; Dreier et al., 2005). This complex is an E3 ubiquitin ligase and is essential for directing proteins to the 26S proteasome for ubiquitin-mediated degradation (Thompson et al., 2021). SCF complexes play critical roles in regulating signaling and regulatory factors and in maintaining proper cellular function (Xie et al., 2019). LIN-23 is required for development and was shown to regulate intestinal development by targeting CDC-25.1/2 for ubiquitination, a process necessary for cell cycle progression (Son et al., 2016). It has also been implicated in the regulation of axon outgrowth in specific classes of motor and sensory neurons (Mehta et al., 2004). In addition, a role for LIN-23 in SKN-1 regulation was reported; LIN-23 negatively regulates SKN-1 during embryonic differentiation (Du et al., 2015). Interestingly, the mammalian ortholog of LIN-23, β-transducin repeat–containing protein (β-TrCP), similarly functions as a negative regulator of Nrf2 by directly targeting it for degradation, thereby modulating the cellular antioxidant response (Chowdhry et al., 2013; Rada et al., 2011). In contrast to these previous findings showing that LIN-23 and orthologs *negatively* regulate SKN-1 and orthologs, we report that LIN-23 *positively* influences SKN-1 activation during pathogen exposure in *C. elegans*, suggesting a context-dependent regulatory relationship.

The goal of this study was to elucidate how LIN-23 promotes SKN-1 activity during pathogen infection. We found that loss of *lin-23* reduces both SKN-1 activity and *C. elegans* survival upon pathogen exposure. We further investigated whether LIN-23 modulates SKN-1 through the p38 MAPK signaling cascade. The data indicate that this regulation occurs downstream of SKN-1 activation, as LIN-23 does not affect SKN-1 phosphorylation or nuclear localization. Furthermore, we present evidence that LIN-23 targets the negative regulator WDR-23 to facilitate SKN-1 activation. In conclusion, genetic analyses support a model in which LIN-23 functions within the WDR-23/SKN-1 regulatory pathway.

## MATERIALS AND METHODS

### *C. elegans* maintenance

*C. elegans* and bacterial strains used in this study are found in Supplementary Table 1. Maintenance of *C. elegans* strains is as previously described (Hope, 1999). Briefly, to generate maintenance plates, the *E. coli* strain OP50 was taken from glycerol stocks and grown on a Luria-Bertani (LB) plate at 37°C overnight. A bacterial sample was then cultured overnight in liquid LB medium at 37°C at 200 RPM. 500 μL of OP50 liquid culture was then seeded onto 100 mm nematode growth (NG) plates and grown overnight at 37°C. Worms were cultured on NG plates with OP50 at 20°C.

### Generation of *C. elegans* strains

The tissue-specific WDR-23B transgenic strain was generated by InVivo Biosystems (IVB) (https://invivobiosystems.com/) using the MosSCI method. Briefly, IVB determined the genomic target site for transgene insertion, selected Mos1 alleles, and verified the presence of the Mos1 transposase. A plasmid construct for transgene expression was designed to express the WDR-23B isoform under the intestinal promoter *vha-6* and tagged with a red fluorescent protein. The plasmid was injected into animals via microinjection, and animals were screened for the transgene, followed by confirmation of its integration. The resulting strain COP2949 was verified by PCR sequencing, and the primers are listed in Supplementary Table 2. Note that a second strain expressing fluorescently tagged WDR-23B under its native promoter was also generated, resulting in strain COP2958. Both strains were then outcrossed to the *lin-23(ot1)* mutant, and the presence of the transgene and mutation was verified via single worm PCR.

### RNA Interference

*E. coli* HT115(DE3) strains containing RNAi constructs of interest (sourced from Geneservices, United Kingdom) were utilized in RNAi experiments. RNAi bacteria were cultured to stationary phase, then 1 mM β-D-1-thiogalactopyranoside (IPTG) was added to the culture and incubated for an additional hour. Following this, the culture was concentrated 5-fold, seeded onto NG plates containing 1 mM IPTG and 100 μg/mL carbenicillin, and incubated at 37°C to induce dsRNA expression. For double RNAi knockdowns, 2 RNAi cultures were mixed at a 1:1 ratio; for all RNAi experiments, the empty vector L4440 in the same parent strain (HT115(DE3)) was used as a control. For RNAi treatment, L1 animals were washed off NG plates, filtered (using a pluriSelect pluriStrainer 10 µm), collected by centrifugation (at 1500 RPM for 1 min), transferred to RNAi-seeded plates, and incubated at 15°C. L1 animals fed on the RNAi bacteria corresponding to the gene to be knocked down until they reached the L4 stage (∼48 hours).

### Bacterial Infection for Microscopy

In this study, animals were infected with either *E. faecalis* (OG1RF) or *P. aeruginosa* (PA14). Synchronized animals at the L4 or young adult stage were exposed to the OG1RF strain for 16 hours on brain heart infusion (BHI) plates, or the PA14 strain for 7 hours on NG plates; *E. coli* on NG plates served as the nonpathogen control. Infections were performed at 25°C, and all infection experiments were performed at least 3 times independently.

### Fluorescence quantification (Confocal microscopy and imaging plate reader)

*Pgst-4*::GFP, *Pgcs-1*::GFP, and SKN-1B/C::GFP expression was assessed using the Olympus FLUOVIEW FV3000 confocal microscope and Fluoview FV315-Sw software. Following bacterial infection or exposure, animals were paralyzed with 25 mM Tetramisole and mounted on a 2% agarose pad for visualization. For *Pgst-4*::GFP and *Pgcs-1*::GFP measurements, the mean level of GFP fluorescence intensity for each animal was quantified using a BioTek Cytation 5 imaging plate reader. For quantification, approximately 50 animals in 100 μL of M9 supplemented with 25 mM of Tetramisole were transferred into a 96-well plate (Corning Incorporated Costar, 3603) and analyzed using Gen5 3.08 software. 3 technical replicates per condition were measured for each biological replicate, and at least 3 biological replicates were performed for each experiment. For detection of SKN-1B/C::GFP expression, a combination of mCherry and EGFP filter sets was used to discriminate intestinal autofluorescence by creating a yellow to red signal. SKN-1B/C::GFP nuclear localization in the intestinal cells was scored as previously described (An and Blackwell 2003; Inoue et al. 2005). Briefly, no nuclear localization, nuclear localization in the anterior or posterior ends of the worm, and nuclear localization throughout all intestinal cells are categorized as low, medium, and high, respectively. Fluorescence microscopy experiments shown were independently repeated at least 3 times.

### Killing assays

Killing assays were performed as previously described (Tan et al. 1999; Garsin et al. 2001). Briefly, 10 µL of OG1RF grown to late log phase was seeded on 35 mm BHI agar plates containing 50 µg/mL gentamycin and 10 µg/mL nystatin and incubated at 37°C for 24 hrs. Palmitic acid (10 µg/mL in ethanol) was used to create a physical barrier on all killing assay plates to prevent animals from escaping. Approximately 90-104 L4 animals were transferred to 3 replica plates and were scored as live or dead at 24-hour intervals beginning on day 3 of infection, and scored until an LD_50_ was reached.

### RNA isolation and qRT-PCR analysis

Approximately 1000 animals were infected with *E. faecalis* OG1RF for 16 hrs or exposed to *E. coli* HTT15 for the same time point; following bacterial exposure, the animals were collected for RNA extraction using Direct-zol RNA Miniprep plus (Zymo Research). cDNA synthesis was performed using HiScript III All-in-one RT SuperMix Perfect for qPCR (Vazyme cat# R333-01) and qRT-PCR was performed using miRNA Unimodal SYBR qPCR Master Mix (Vazyme cat# MQ102-02) in a Bio-Rad CFX 96 Real-Time system. Primers sequences used can be found in Supplementary Table 2. All values were normalized to *act-1*.

### Single worm PCR

The *lin-23(ot1)* strain was backcrossed 4x into our laboratory’s N2 strain. The backcrossed mutant was then crossed into various fluorescent reporter strains (listed in Supplementary Table 1). To detect the *lin-23(ot1)* mutation, single worm PCRs were performed, followed by a restriction enzyme test. To collect DNA from single worms, individual adult-stage animals were placed into 12 μL of worm lysis buffer and frozen at – 80°C for 1 hr. Then, the animals were heated using the thermal cycles 65°C for 1 hr followed by 95°C for 15 mins. 4 μL of single worm DNA was used in conventional PCR. The primers used are listed in Supplementary Table 2. The PCR was performed using the following parameters: initial step at 95°C for 5 min, followed by 34 cycles of 95°C for 30 s, 60°C for 40 s, 72°C for 1.5 min, and a final extension cycle at 72°C for 5 min. Following amplification, DNA was digested with the HaeIII restriction enzyme according to the NEB protocol. The digested DNA was then mixed with gel loading dye and loaded into an agarose gel. Visualization and quantification of the bands of interest were performed using a 1.5% agarose gel and GelGreen nucleic acid stain using a ChemiDoc MP Imaging System (Bio-Rad).

### Western Blot Analysis

Approximately 3,000 animals were collected following *E. faecalis* OG1RF exposure, washed, and boiled in the sample buffer. Following centrifugation, the supernatant was collected, flash-frozen, and then boiled at 80°C for 10 minutes. Protein lysates were resolved on a SDS-polyacrylamide gel via electrophoresis, transferred to a PVDF membrane, and incubated with a rabbit anti-phospho-p38 MAPK (Cell Signaling, #9211) using a 1:2000 dilution. An identical membrane was incubated with mouse anti-tubulin (Sigma, #T9026) at a 1:4000 dilution for a loading control. For secondary antibodies, blots were incubated with 1:2000 secondary HRP-conjugated anti-mouse (for anti-tubulin) and anti-rabbit (for anti-phospho-p38) antibody for 1 hour and subsequently washed 3 times for 5-min intervals. 1 milliliter of SuperSignal West Atto Ultimate Sensitivity Substrate (Thermo Fisher, A38555) was added to each membrane and they were visualized using a ChemiDoc MP Imaging System (Bio-Rad). ImageJ software was used for quantifying the immunoblot.

### Sodium arsenite assays

To test the effects of arsenite stress on the *lin-23(ot1)* mutant, synchronized L1 animals were exposed to RNAi as follows: *lin-23* mutants were exposed to *cdc-25* RNAi, and N2 and *skn-1(zu135)* animals were exposed to *lin-23* and *cdc-25* RNAi. All animals were incubated at 15°C until the L4 stage was reached. Following exposure to RNAi, the animals were washed and transferred to a 35 mm NG plate supplemented with 10 mM of arsenite (NaAsO_2_) and seeded with *E. coli* OP50. Approximately 30 animals per condition were transferred to a plate (in triplicate) and incubated at 20°C for 8 hours, with live/dead scoring being performed at the end of each hour. For qRT-PCR analysis, approximately 1000 animals per RNAi condition were exposed to sodium arsenite for 4 hours, followed by RNA extraction.

### Statistical analyses

GraphPad Prism 11 was used for data analysis. Following categorical scoring of confocal images, statistical significance was determined by the chi-square of Fisher’s exact tests. For quantification of fluorescent measures, qRT-PCR/RT-PCR, and western blots, significance was determined using unpaired t tests for experiments with only two samples, or one-way ANOVA followed by Dunnett’s multiple comparison tests for experiments with multiple samples. Mantel-Cox log-rank analysis was used to compare survival curves and to calculate the median survival. Comparisons of interest and their statistical significance are indicated in the figures. For all statistical tests, P values < 0.05 were considered statistically significant. *P <0.05, **P < 0.01, ***P < 0.001, ****P < 0.0001.

## RESULTS

### Loss of LIN-23 decreases SKN-1 activity during pathogen exposure

During infection of *C. elegans* with either *E. faecalis* or *P. aeruginosa,* there is an accumulation of the bacteria in the intestine and production of phase II detoxification enzymes encoded by genes directly regulated by SKN-1 (Hoeven et al., 2011; Papp et al., 2012). Reporters for two of these genes, *Pgst-4*::GFP and *Pgcs-1*::GFP, were used to identify factors that, when reduced by RNAi, prevented their activation by *E. faecalis* and *P. aeruginosa*, respectively. As shown in **Fig. 1a-b**, loss of *lin-23* by RNAi reduced the expression of these reporters in the intestine of *C. elegans* in a manner comparable to the positive control, *skn-1* RNAi. The phenotypes were visible in both representative images and by quantifying the fluorescent signal with an imaging plate reader (**Fig. 1c-d**).

**Fig. 1.**
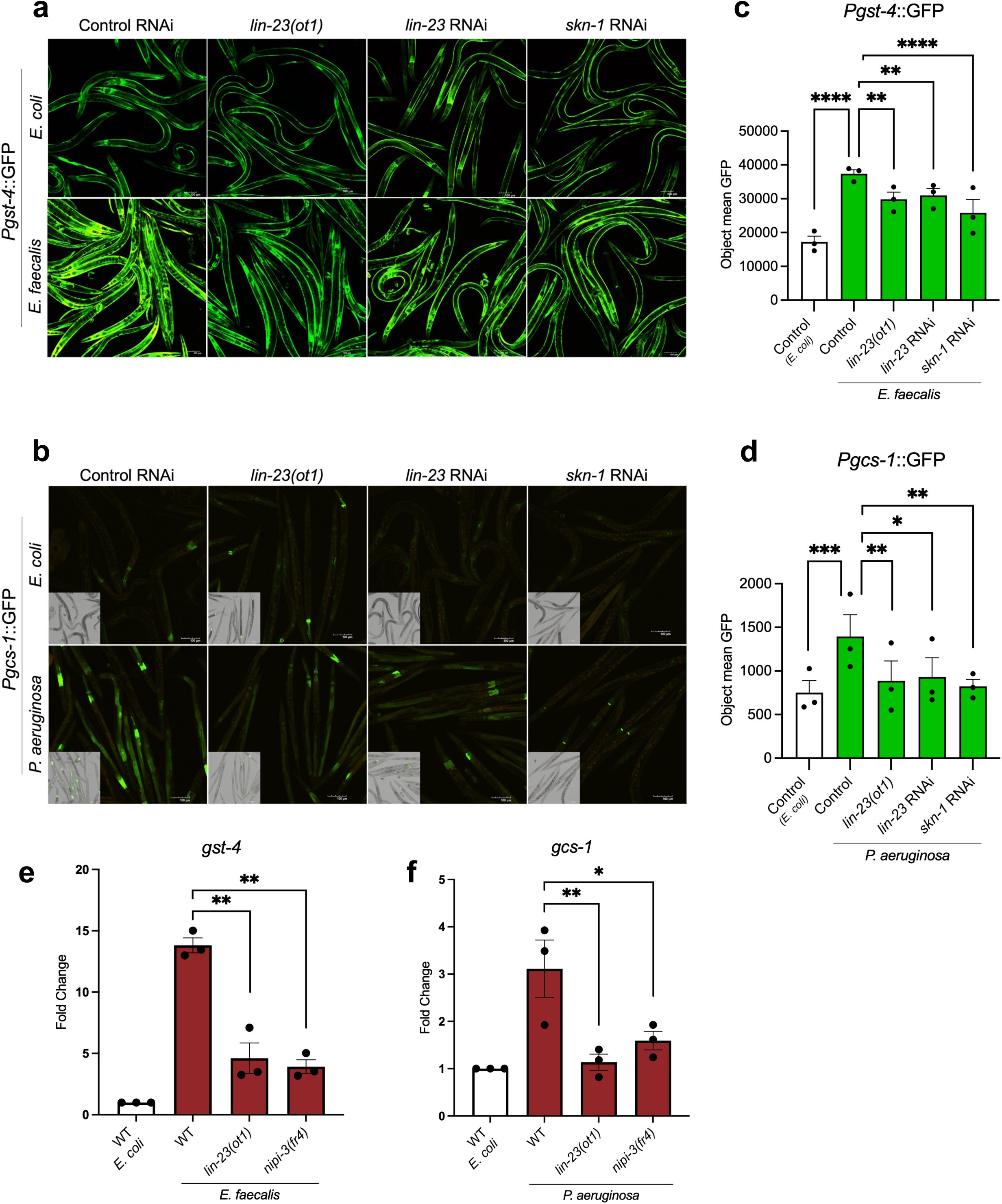
Loss of LIN-23 decreases SKN-1 activity during pathogen exposure. a) and b) Expression patterns in *Pgst-4::GFP* and *Pgcs-1::GFP* tagged worms following exposure to *E. faecalis*, *P. aeruginosa*, or *E. coli* following RNAi knockdown of the indicated genes and in a *lin-23(ot1)* background. Images include 100 µm scale bars. c) and d) Quantification of GFP fluorescence in animals with *Pgst-4::GFP* and *Pgcs-1::GFP* expression reporters, as shown in panels a) and b), was performed using an imaging plate reader (BioTek Cytation 5). The *y*-axis indicates the average pixel count measured in worms following pathogen exposure. Analysis was conducted on three biological replicates, each with approximately 50 worms. Error bars indicate the standard error of the mean (SEM). Asterisks indicate the statistical significance of the bracketed comparisons. \**P* < 0.05; \*\**P* < 0.01; \*\*\**P* < 0.001; \*\*\*\**P* < 0.0001; ns, not significant. e) and f) *gst-4* and *gcs-1* expression levels were measured by qRT-PCR in N2 wild-type, *lin-23(ot1)*, and *nipi-3(fr4)* mutant worms exposed to *E. faecalis*, *P. aeruginosa* or *E. coli*. Gene expression values were normalized to the housekeeping gene, *act-1*, and are relative to the wild-type worms exposed to *E. coli* and set to 1. Analysis was conducted on three biological replicates; error bars represent the SEM of the biological replicates, and asterisks indicate the statistical significance of the bracketed comparisons.

To further investigate and confirm this phenotype, we utilized a *lin-23* mutant strain. Due to LIN-23’s essentiality in regulating postembryonic cell divisions, it is not possible to study a null mutant. However, a missense mutant, *lin-23(ot1)*, was previously identified, in which a single amino acid change, from proline 610 to serine, occurs near the C-terminus. This mutation is predicted to affect the protein-binding motif PAPP, which is vital for the function of F-box proteins as the substrate-binding subunit within the SCF complex (Mehta et al., 2004). The mutant was obtained and crossed to the strains containing the SKN-1-regulated fluorescent reporters. Both *Pgst-4*::GFP and *Pgcs-1*::GFP displayed reduced expression during infection, as shown in the representative images in **Fig. 1a and b**. The decrease in reporter expression was comparable to that observed in the *lin-23* and *skn-1* RNAi conditions (**Fig. 1c-d**). To corroborate these results, the expression levels of the endogenous genes, *gst-4* and *gcs-1,* were measured in the LIN-23 mutant; expression of both genes was significantly decreased during infection compared to the wild-type N2 strain (**Fig. 1e and 1f**). A NIPI-3 mutant served as a positive control, as previous work demonstrated that loss of NIPI-3 results in a significant reduction in *gst-4* and *gcs-1* during infection (C. Wu et al., 2021).

### Loss of LIN-23 decreases *C. elegans* survival on pathogen

Because loss of SKN-1 activity reduces pathogen resistance, we also assessed the role of LIN-23 in survival following pathogen infection. Upon exposure to *E. faecalis*, both LIN-23 mutant animals and animals exposed to LIN-23 RNAi exhibited sensitivity to the pathogen as compared to control RNAi and the wild-type strain, respectively (**Fig. 2a**). An additional pathogen resistance assay was conducted using a loss-of-function allele of *skn-1*, shown in **Fig. 2b**. As expected, these mutants exhibited increased sensitivity to pathogen infection compared to wild-type animals. Upon exposure to *lin-23* RNAi, no significant change in susceptibility was observed in the *skn-1* mutants. This lack of an additive effect suggests that *lin-23* and *skn-1* function within the same genetic pathway. Together, these findings support a role for LIN-23 in SKN-1–mediated pathogen resistance.

**Fig. 2.**
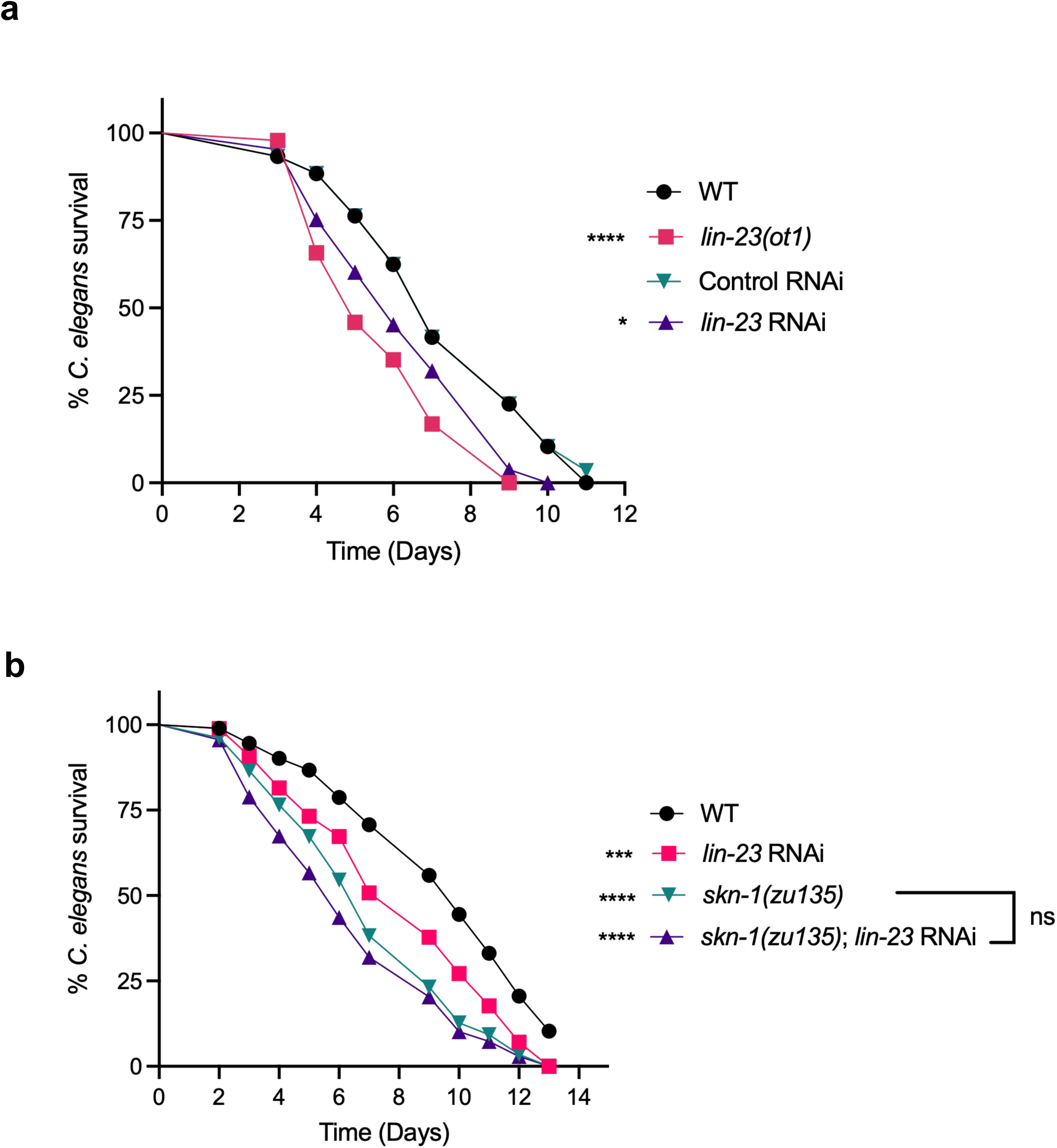
Loss of LIN-23 decreases *C. elegans* survival during *E. faecalis* infection. a) Survival curves of wild-type and *lin-23(ot1)* animals exposed to control or *lin-23* RNAi during infection with *E. faecalis*. Asterisks indicate statistical significance for comparisons between wild-type and *lin-23(ot1)* animals, as well as between control RNAi and *lin-23* RNAi conditions. b) Survival curves of wild-type and *skn-1(zu135)* animals following exposure to *lin-23* RNAi during infection with *E. faecalis*. Asterisks indicate statistical significance relative to wild-type animals. For all survival assays, worms were also treated with *cdc-25.1* RNAi to induce sterility (Hoeven et al., 2011). Graphs shown in both panels are representative of three independent trials. Sample sizes, median survival, and *P*-values of all trials are given in Supplementary Table 3. \**P* < 0.05; \*\*\**P* < 0.001; \*\*\*\**P* < 0.0001; ns, not significant.

### LIN-23 also modulates survival following exposure to NaAsO_2_

As previously reported, LIN-23 negatively regulates SKN-1 during embryogenesis, thereby controlling the progression of differentiation (Du et al., 2015). In contrast, our findings indicate that LIN-23 functions as a positive regulator of SKN-1 during pathogen infection. Given that distinct stressors can activate different regulatory pathways, we next investigated whether LIN-23 also modulates SKN-1 activity in response to oxidative stress induced by chemical agents. To test this, we exposed *lin-23(ot1)* mutants to sodium arsenite (NaAsO₂) and observed increased sensitivity compared to wild-type animals (**Fig. 3a**). Furthermore, *skn-1* mutants treated with *lin-23* RNAi exhibited a level of sensitivity to sodium arsenite comparable to that of untreated *skn-1* mutants, consistent with dependence on SKN-1 under these conditions. To further examine the dependence on SKN-1, qRT-PCR analysis was performed to measure expression of the SKN-1-dependent genes *gst-4* and *gcs-1* in sodium arsenite-treated animals, as shown in **Fig. 3b-c**. *skn-1* animals exhibited significantly reduced expression of these genes. Animals with reduced LIN-23 levels, either by mutation or RNAi, showed a trend towards reduction; however, the results did not reach statistical significance. Overall, the results show that LIN-23 modulates stress resistance in oxidant-exposed animals, and suggest that the effect is partially dependent on SKN-1.

**Fig. 3.**
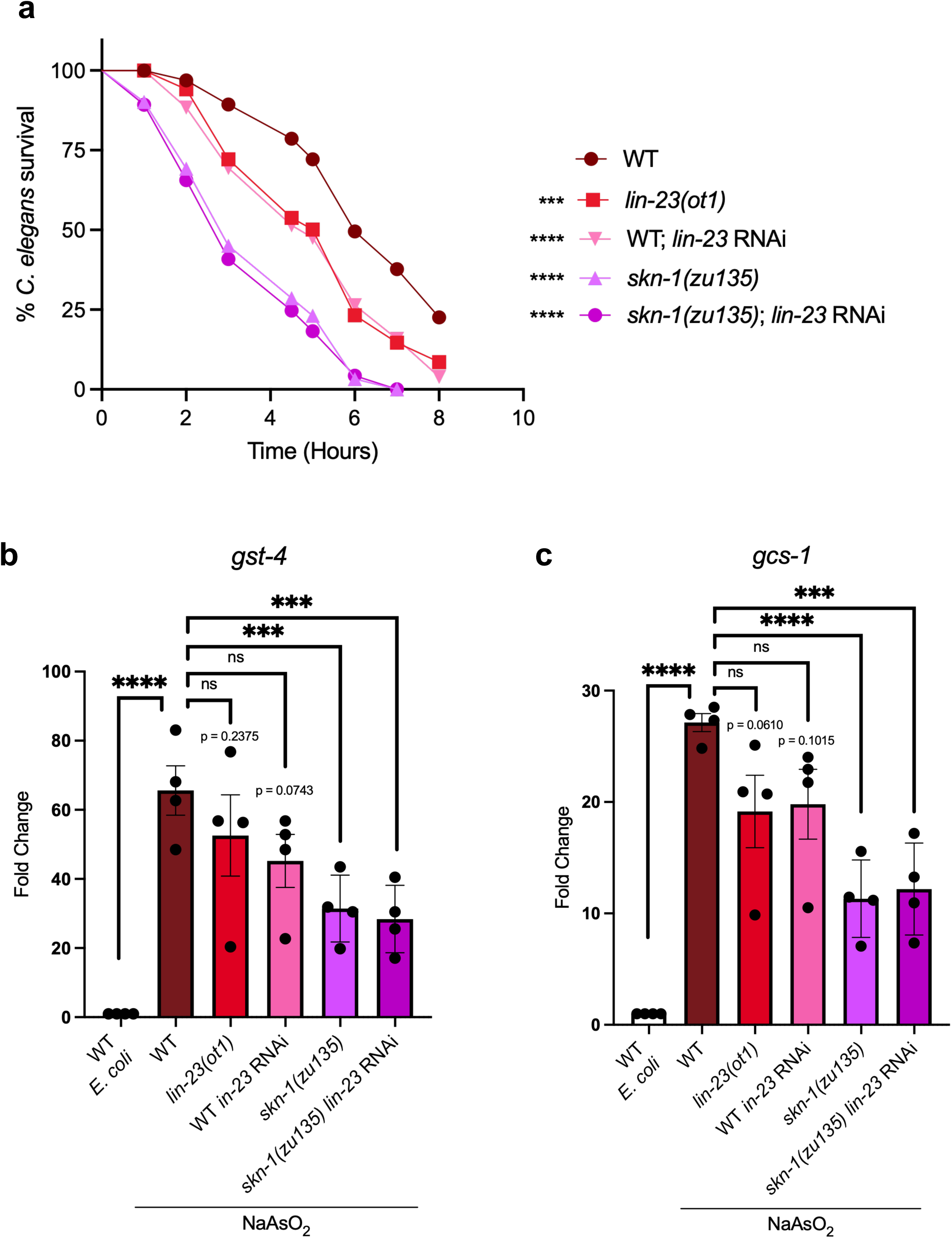
LIN-23 affects survival following sodium arsenite exposure. a) Survival curves of N2 wild-type, *lin-23(ot1)*, and *skn-1(zu135)* animals treated or untreated with *lin-23* RNAi and exposed to sodium arsenite (NaAsO_2_). Asterisks indicate statistical significance relative to N2 wild-type animals. For all survival assays, worms were also treated with *cdc-25.1* RNAi. \*\*\**P* < 0.001; \*\*\*\**P* < 0.0001. The data shown is representative of three independent trials. Sample sizes, median survival, and *P*-values of all trials are given in Supplementary Table 3. b) and c) *gst-4* and *gcs-1* expression levels were measured by qRT-PCR in N2 wild-type, *lin-23(ot1)*, and *skn-1(zu135)* mutant worms exposed to *E. coli* on plates supplemented with and without sodium arsenite. Gene expression values were normalized to the housekeeping gene, *act-1*, and are relative to the wild-type worms not exposed to sodium arsenite and set to one. Analysis was conducted on four biological replicates; error bars represent the SEM of the biological replicates, and asterisks indicate the statistical significance of the bracketed comparisons. \**P* < 0.05; \*\**P* < 0.01; \*\*\**P* < 0.001; \*\*\*\**P* < 0.0001; ns, not significant.

### LIN-23 regulates SKN-1 downstream of the p38 MAPK pathway

During pathogen infection, host-generated reactive oxygen species (ROS) trigger the p38 MAPK signaling cascade, resulting in the phosphorylation of SKN-1 via the mitogen-activated protein kinase PMK-1. In turn, phosphorylated SKN-1 localizes to the nuclei of intestinal cells, where it is activated and aids in the transcription of phase II detoxification enzymes (Hoeven et al., 2011). To elucidate the mechanisms by which LIN-23 regulates SKN-1, we examined p38 MAPK pathway activation using PMK-1 phosphorylation levels as a readout. As shown in **Fig. 4a**, neither the *lin-23* mutant nor animals exposed to *lin-23* RNAi exhibited reduced levels of phosphorylated PMK-1 following infection with *E. faecalis*. Notably, the levels were slightly elevated in the *lin-23* mutant (**Fig. 4b**). This could be due to a block in activation downstream of the p38 MAPK pathway, preventing negative feedback regulation on the phosphorylation cascade. Overall, these results show that LIN-23 does not negatively impact the phosphorylation of the p38 MAPK, PMK-1, which is critical for SKN-1 activity.

**Fig. 4.**
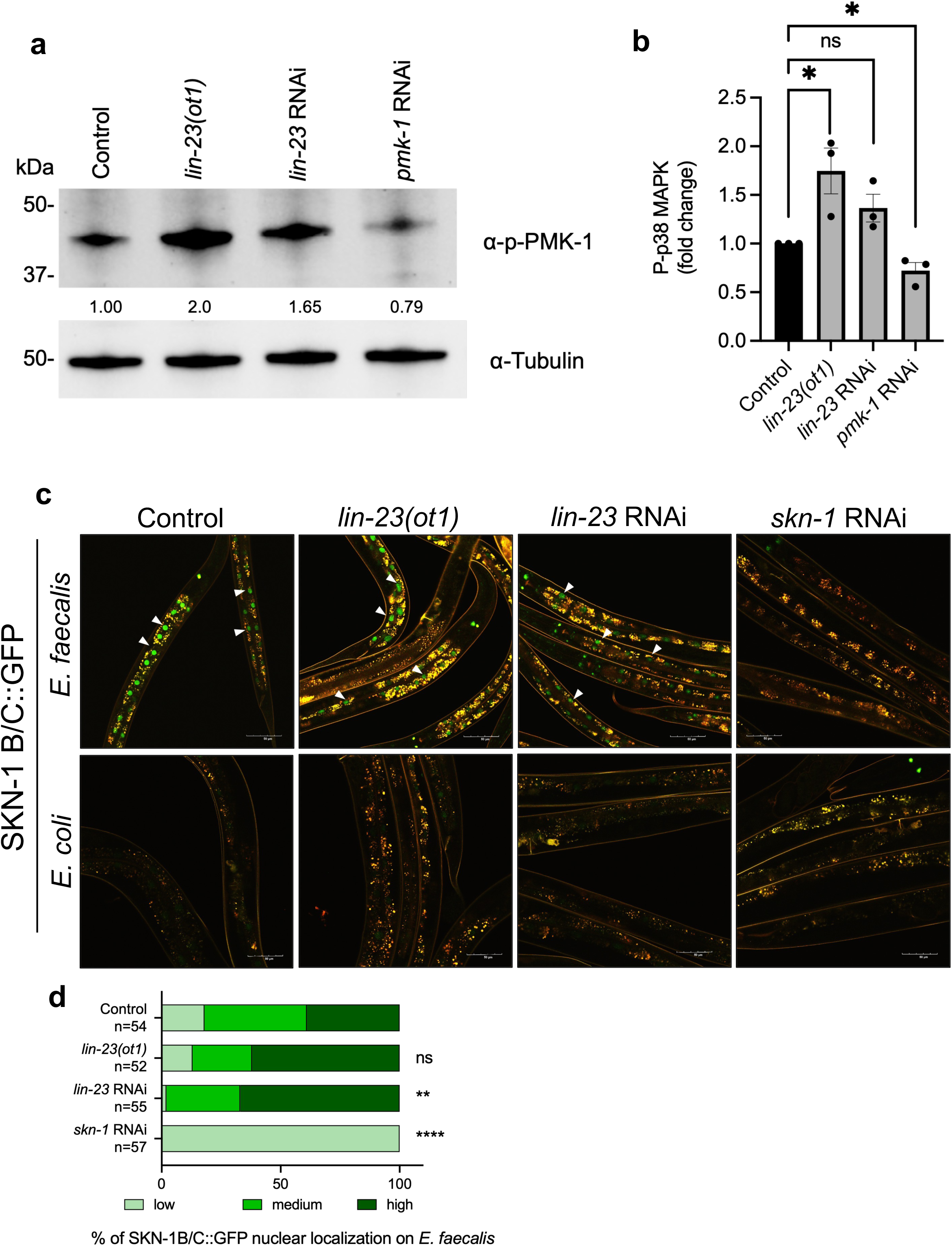
LIN-23 regulates SKN-1 downstream of the p38 MAPK cascade. a) Western blot analysis of PMK-1 phosphorylation levels using an α-phospho-p38 antibody and an α-tubulin antibody (loading control) from lysates of the indicated animals exposed to *E. faecalis*. Blots shown are representative. The average band intensities relative to α-tubulin across 3 biological replicates are presented below. b) Analysis of blots measuring band intensities. Bands were densitometrically analyzed using ImageJ software, and, following normalization to the housekeeping gene, tubulin, the levels of p-PMK-1 were calculated with the control condition set to one. Error bars represent the SEM of the averaged biological replicates. The asterisks indicate the statistical significance of the bracketed comparisons. \**P* < 0.05; ns, not significant. c) Wild-type and *lin-23(ot1)* worms containing the integrated SKN-1B/C::GFP transgene were exposed to *E. faecalis* or *E. coli*. SKN-1B/C::GFP localization was observed by fluorescence microscopy. Examples of intestinal nuclei displaying clear nuclear localization are indicated by white arrows. Scale bars represent 50 μm. d) SKN-1C nuclear localization was scored based on the GFP signal (low, medium, high), and the percentage for each category was quantified. The number of worms used in scoring each experimental condition is indicated (*n*). Levels of SKN-1B/C::GFP nuclear localization in the intestines of animals on pathogen in the *lin-23(ot1)* mutants or wild-type worms following exposure to *lin-23* and *skn-1* RNAi were compared to the control. Asterisks indicate the statistical significance of the bracketed comparisons. \*\**P* < 0.01; \*\*\*\**P* < 0.0001; ns, not significant.

We next investigated whether LIN-23 affects SKN-1 nuclear localization in the intestine. To do this, a SKN-1 B/C::GFP fluorescent reporter strain was crossed to the *lin-23* mutant and subsequently infected with *E. faecalis*. During infection, loss of *lin-23,* either by mutation or by RNAi, did not reduce SKN-1 nuclear localization, as shown in the representative images in **Fig. 4c** compared with control animals. *skn-1* RNAi served as a positive control for loss of visible SKN-1 nuclear localization. Quantification of the visibility of intestinal nuclei was performed using categorical scoring (see Methods (Gabaldón et al., 2024; C. Wu et al., 2021)) and confirmed the qualitative observations (**Fig. 4d**). As exposure to *P. aeruginosa* also activates SKN-1 nuclear localization (Papp et al., 2012), we also examined animals exposed to this pathogen. The observed phenotype, loss of *lin-23* NOT affecting SKN-1 localization, was also apparent during infection with *P. aeruginosa* (Supplemental Fig. 1). Together, these results suggest LIN-23 acts downstream of the p38 MAPK pathway and SKN-1 nuclear localization to modulate SKN-1 activity during infection.

### *lin-23* functions within the *wdr-23/skn-1* genetic pathway

Because of its role as an F-box protein *positively* regulating SKN-1 activity, a plausible hypothesis is that LIN-23 is targeting a negative regulator of SKN-1 for degradation. Therefore, we investigated whether there is a genetic interaction with WDR-23, a well-known negative regulator of SKN-1 (Choe et al., 2009; Spatola et al., 2019; Tang & Choe, 2015). To determine whether *lin-23* functions within the same genetic pathway as *wdr-23*, we performed an epistasis analysis using a *wdr-23* null mutant carrying a SKN-1–dependent fluorescent reporter, combined with RNAi treatment. As shown in **Fig. 5a**, we observed very high levels of reporter activation in the *wdr-23* background under both non-pathogenic and pathogenic conditions, as had been previously reported (Choe et al., 2009; Tang & Choe, 2015). However, no change in reporter fluorescence was observed between control and *lin-23* RNAi, consistent with WDR-23 acting downstream of LIN-23 in the same genetic pathway. As expected, *skn-1* RNAi significantly reduced fluorescence, serving as a positive control. Quantification of fluorescence using an imaging plate reader confirmed these observations under both conditions (**Fig. 5b–c**).

**Fig. 5.**
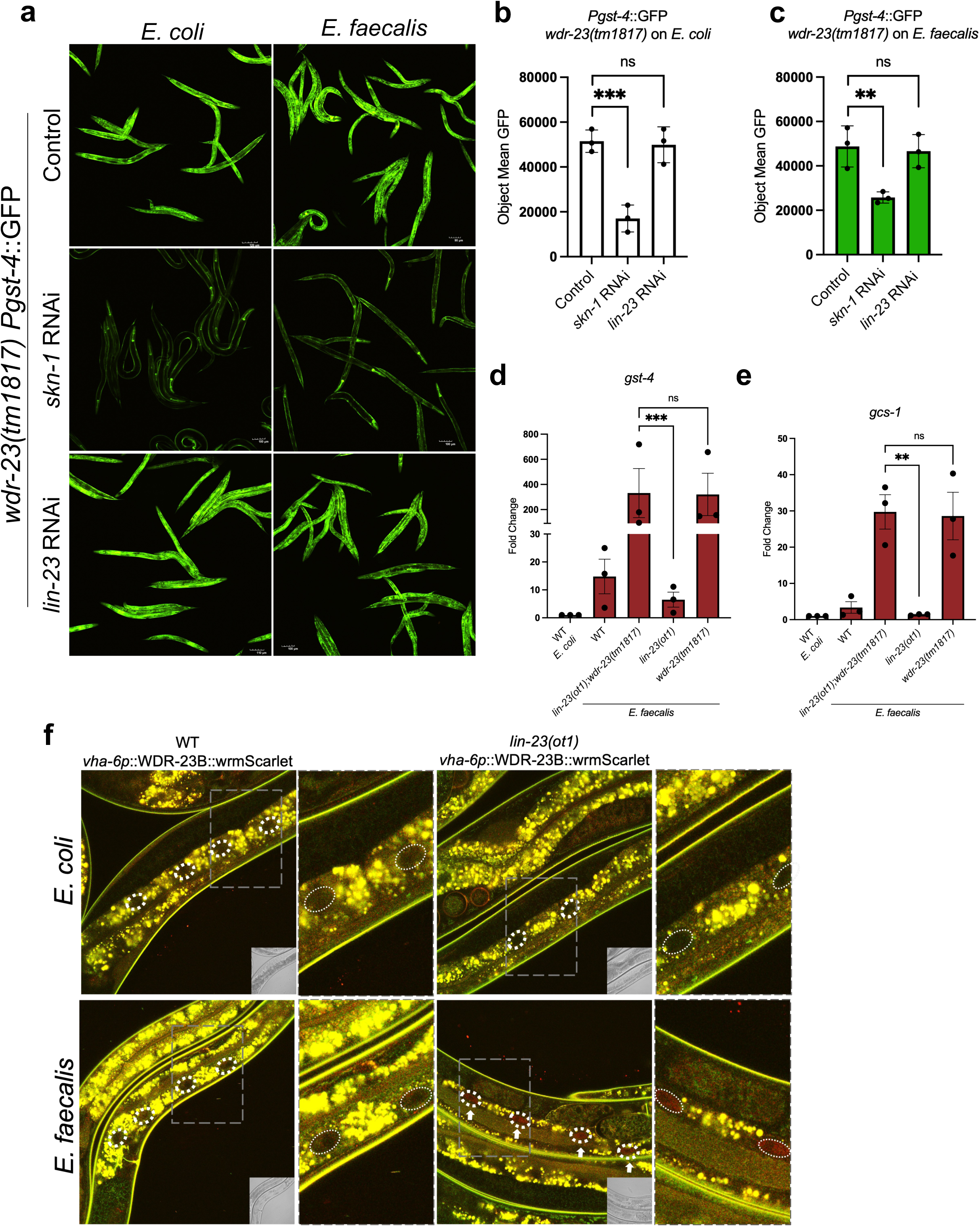
*lin-23* functions within the *wdr-23/skn-1* genetic pathway. a) Expression patterns of *Pgst-4::GFP* in *wdr-23(tm1817)* worms exposed to *E. coli* or *E. faecalis* following RNAi knockdown of the indicated genes. b) and c) Quantification of *Pgst-4::GFP* fluorescence in *wdr-23(tm1817)* animals exposed to *E. coli* or *E. faecalis*, as shown in panel a), was performed using an imaging plate reader (BioTek Cytation 5). The *y*-axis indicates the average pixel count measured in worms. Analysis was conducted on three biological replicates, each with approximately 50 worms. Error bars indicate the standard error of the mean (SEM). Asterisks indicate the statistical significance of the bracketed comparisons. \*\**P* < 0.01; \*\*\**P* < 0.001; ns, not significant. d) and e) Endogenous *gst-4* and *gcs-1* expression levels were measured by qRT-PCR in N2 wild-type, *lin-23(ot1)*, and *wdr-23(tm1817),* and *lin-23(ot1);wdr-23(tm1817)* mutant worms exposed to *E. faecalis* or *E. coli*. Gene expression values were normalized to the housekeeping gene, *act-1*, and are relative to the wild-type worms exposed to *E. coli* and set to one. Analysis was conducted on three biological replicates; error bars represent the SEM of the biological replicates, and asterisks indicate the statistical significance of the bracketed comparisons. \*\**P* < 0.01; \*\*\**P* < 0.001; ns, not significant. f) WDR-23B::wrmScarlet expression under the intestinal *vha-6* promoter in wild-type and *lin-23(ot1)* backgrounds following exposure to *E. coli* or *E. faecalis*. Presence and absence of nuclear WDR-23B::wrmScarlet expression in the *lin-23(ot1)* background following infection with *E. faecalis* is shown by white dashed lines.

To further corroborate the genetic interaction between *lin-23* and *wdr-23*, we performed an additional epistasis analysis to assess SKN-1 activity in *lin-23, wdr-23,* and *lin-23;wdr-23* double mutants. We measured endogenous expression of the SKN-1 target genes *gst-4* and *gcs-1* by qRT-PCR during infection (**Fig. 5d–e**). These results showed increased SKN-1 activity in the *wdr-23* mutant, whereas the *lin-23* mutant had decreased SKN-1 activity, as expected. The *wdr-23* mutant was crossed with the *lin-23* mutant, and the resulting double mutant exhibited high endogenous expression of *gst-4* and *gcs-1*, comparable to the *wdr-23* single mutant (**Fig. 5d–e**). Because the double mutant phenocopied the *wdr-23* mutant rather than the *lin-23* mutant, these data further support a model in which *wdr-23* acts downstream of *lin-23* to regulate SKN-1 activity.

To investigate if LIN-23 affects WDR-23 levels, consistent with LIN-23 targeting WDR-23 for degradation, we collaborated with InVivo Biosystems, to design and generate transgenic *C. elegans* strains in which WDR-23B is tagged with wrmScarlet under an intestinal-specific promoter (*vha-6*) and its native promoter. Recall that this isoform had previously been characterized as being expressed in the nuclei and able to complement the WDR-23 stress-related phenotypes (Lo et al., 2017; Staab et al., 2013; Spatola et al., 2019). Both transgenic strains were crossed to the *lin-23*(*ot1*) strain. When examining the strains by fluorescence microscopy under both non-infectious and infectious conditions, WDR-23B expression driven by its native promoter was not detected in either the wild-type or *lin-23(ot1)* strain (Supplemental Fig. 2). This may be due to low endogenous expression, as we generated these strains using the MosSCI technique rather than an overexpression construct, as was done previously (Staab et al., 2013). We also did not detect any WDR-23B expression in either strain under the intestinal promoter during non-infectious conditions (**Fig. 5f**). However, following exposure to *E. faecalis,* WDR-23B expression was detected, but only in the *lin-23(ot1)* strain background. Specifically, we observed WDR-23B localized to the intestinal nuclei (outlined in white), consistent with previous studies showing nuclear localization of the B isoform (**Fig. 5f**). This increase in detectable WDR-23B during infection in the absence of functional LIN-23 supports a model in which LIN-23 targets WDR-23B for degradation during infection to promote SKN-1 activity.

## DISCUSSION

Thus far, LIN-23 has been reported to regulate cell cycle progression (Kipreos et al., 2000; Son et al., 2016), axon outgrowth (Mehta et al., 2004), and synaptic transmission (Dreier et al., 2005). Furthermore, LIN-23 has been reported to negatively regulate SKN-1 during embryogenesis as part of an E3 ligase that targets the zinc-finger protein OMA-1 for degradation, thereby promoting SKN-1 turnover and facilitating EMS-to-MS differentiation (Du et al., 2015). In contrast to these regulatory roles during embryogenesis and development, our findings establish a previously unrecognized role for LIN-23 as a positive regulator of SKN-1 in adult animals under stressful conditions. Specifically, loss of LIN-23 resulted in reduced SKN-1 activity, as evidenced by decreased expression of the SKN-1 target genes *gst-4* and *gcs-1* during pathogen infection. Consistent with this, *lin-23* mutants exhibited increased sensitivity to pathogen infection, a phenotype dependent on SKN-1, as demonstrated by pathogen resistance assays.

Interestingly, our results suggest that the regulatory role of LIN-23 on SKN-1 during stress responses in adult animals may be context dependent. While animals with reduced LIN-23 levels exhibited significantly increased sensitivity when exposed to sodium arsenite (**Fig. 3a**), these mutants showed only a slight, not statistically significant, decrease in *gst-4* and *gcs-1* expression (**Fig. 3b-c**). In contrast, loss of LIN-23 resulted in significantly lower *gst-4* and *gcs-1* expression during pathogen exposure (**Fig. 1**). This suggests that the decrease in stress resistance is not fully explained by LIN-23 modulating SKN-1-dependent genes under sodium arsenite stress conditions. It would be of interest to examine the role of LIN-23 and its dependence on SKN-1 under a range of oxidative stress conditions in future studies.

Having established the functional importance of LIN-23 in SKN-1-mediated stress responses, we next investigated the underlying mechanism. First, we examined whether LIN-23 influences the p38 MAPK signaling pathway, a well-characterized activator of SKN-1 during infection. However, loss of LIN-23 did not reduce PMK-1 phosphorylation, suggesting that LIN-23 does not act by inhibiting this pathway. Additionally, we assessed SKN-1 subcellular localization and found that its nuclear localization was not impeded in the absence of LIN-23, indicating that LIN-23 is not required for SKN-1 nuclear localization. Based on this negative data and the proposed role of LIN-23 as a component of an SCF ubiquitin ligase complex, a logical hypothesis was that LIN-23 functions by targeting a negative regulator of SKN-1 for degradation. In this context, we investigated whether LIN-23 operates within the established WDR-23/SKN-1 genetic pathway, given that WDR-23 is a well-characterized negative regulator of SKN-1 during stress responses. Moreover, the nuclear isoform of this protein, WDR-23B, was previously characterized as the one necessary for oxidative stress resistance (Lo et al., 2017; Staab et al., 2013; Spatola et al., 2019). Our epistasis analyses support this model. Specifically, loss of LIN-23 did not reduce SKN-1 activity in a *wdr-23* null mutant background, and furthermore, the *lin-23;wdr-23* double mutant phenocopied the *wdr-23* single mutant rather than the *lin-23* mutant, consistent with *wdr-23* functioning downstream of *lin-23*. Notably, our localization study provides additional support for our hypothesis as examination of intestinal WDR-23B expression revealed that, during infection, WDR-23B is detected in the nuclei in the absence of LIN-23, whereas its signal is diminished when LIN-23 is present. Together, these findings support a model diagrammed in Fig. 6 in which LIN-23 regulates SKN-1 activity by modulating WDR-23, potentially through ubiquitin-mediated turnover.

**Fig. 6.**
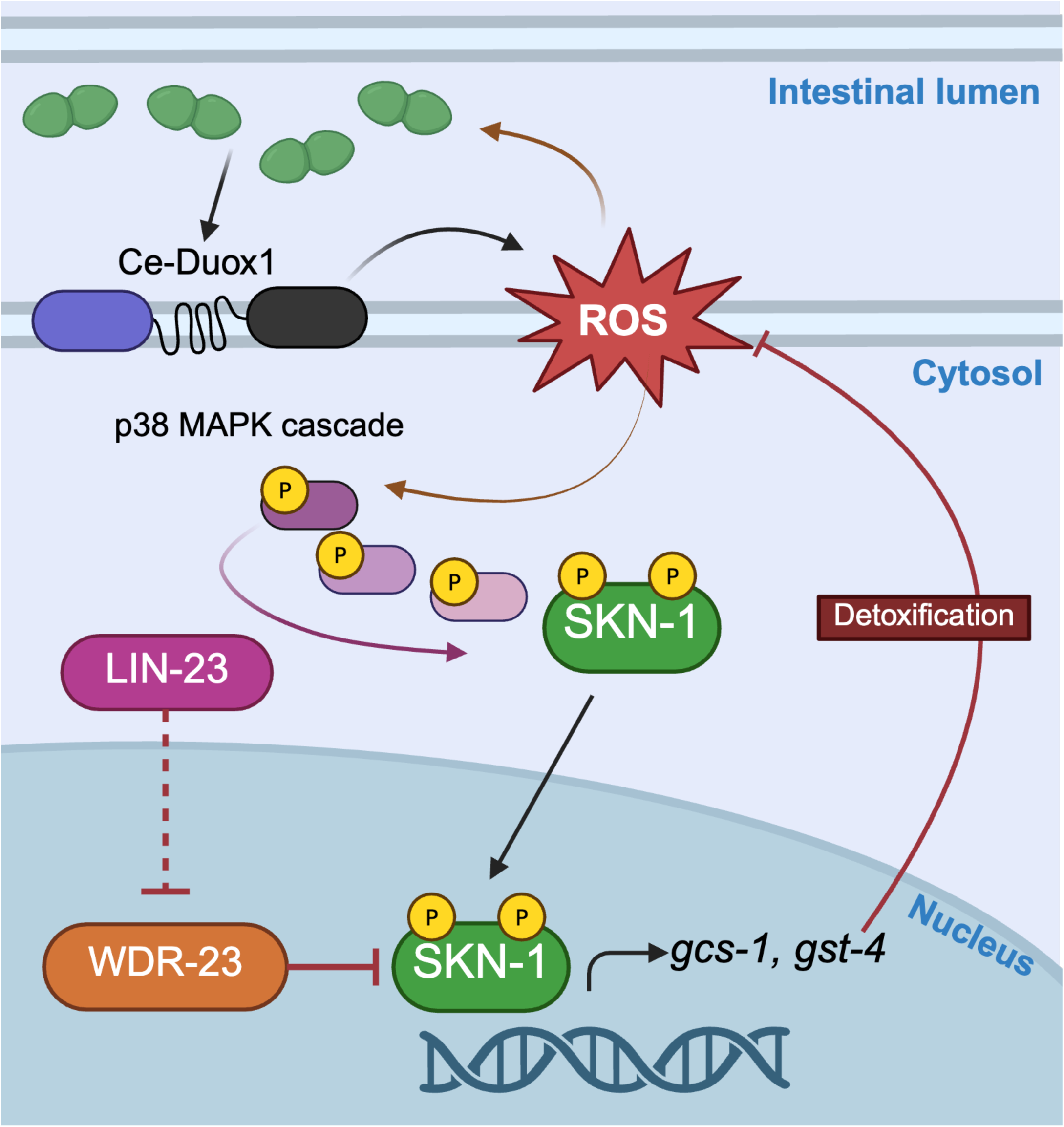
Proposed model of LIN-23-mediated regulation of SKN-1. ROS generated during pathogen infection by Ce-Duox1/BLI-3 activates p38 MAPK signaling and SKN-1 nuclear localization and transcriptional activity as established in previous work (Hoeven et al 2011). Here we show that loss of LIN-23 does not impair p38 MAPK signaling or SKN-1 nuclear localization. Instead it appears to regulate SKN-1 activity by modulating WDR-23, a negative regulator of SKN-1. Specifically, we observe enhanced levels of the nuclear WDR-23B isoform upon loss of LIN-23. Please refer to the main text for details. Diagram created in https://BioRender.com.

This model is appealing given that WDR-23 is a well-established negative regulator of SKN-1, and our findings identify LIN-23 as a positive regulator of SKN-1 in adult animals under pathogen and oxidant stress. Because LIN-23 functions as part of an SCF E3 ubiquitin ligase complex and has primarily been characterized as a negative regulator of its substrates, it is plausible that LIN-23 promotes SKN-1 activity by targeting WDR-23 for degradation. Supporting this idea, WDR-23 has previously been reported to be negatively regulated by another F-box protein, XREP-4 (C.-W. Wu et al., 2017). Notably, several phenotypes described in this study parallel the observations reported here for LIN-23. Specifically, loss of LIN-23, like loss of XREP-4, results in increased sensitivity to stress, but does not suppress the *gst-4p*::GFP constitutive activation observed in a *wdr-23* null mutant, nor affect SKN-1B/C::GFP accumulation in intestinal nuclei. The similarities between the phenotypes associated with XREP-4 and LIN-23 suggest that WDR-23 may be regulated by multiple F-box proteins. Such a model is consistent with the broader complexity of SKN-1 regulation. SKN-1 is a multifunctional transcription factor involved in diverse biological processes. F-box proteins interact with multiple substrates and participate in diverse regulatory networks. Furthermore, the *C. elegans* genome encodes more than 500 F-box proteins. Therefore, it is possible, and in fact likely, that multiple F-box proteins are targeting the same substrate in a context-dependent manner (Skaar et al., 2013).

Although our genetic analyses under adult oxidative stress conditions provide evidence that LIN-23 functions within the same pathway as WDR-23, it remains unknown whether these proteins interact directly. Biochemical assays assessing physical binding between LIN-23 and WDR-23 and examining changes in WDR-23 ubiquitination in the presence or absence of LIN-23 would provide further mechanistic support for a model in which LIN-23 directly regulates WDR-23 within the WDR-23/SKN-1 regulatory pathway, and are underway. However, it remains possible that LIN-23 influences SKN-1 activity through an alternative regulatory factor. F-box proteins can interact with multiple substrates and are capable of mediating diverse cellular outcomes beyond proteasomal degradation (Skaar et al., 2013). Therefore, while our data are consistent with a model in which LIN-23 directly regulates WDR-23, LIN-23 may instead participate in a parallel or upstream pathway that indirectly modulates WDR-23 activity and, consequently, SKN-1 function. To address this possibility, future studies will be required to identify additional LIN-23-interacting cofactors involved in stress response regulation. Importantly, LIN-23 has previously been implicated in regulating development and longevity (Chaudhari & Kipreos, 2017; Maro et al., 2009), processes in which SKN-1 also plays central roles. Taken together, these observations suggest that the relationship between LIN-23 and SKN-1 may be more complex than a direct pathway involving WDR-23 degradation alone and may change depending on the tissue, developmental stage, or stress under which it is assessed.

While we have established a role for LIN-23 in regulating SKN-1 during pathogen stress, the underlying molecular mechanisms remain to be fully elucidated. Our findings suggest that LIN-23 functions independently of the p38 MAPK signaling cascade and instead may regulate SKN-1 through interaction with the negative regulator WDR-23. However, additional studies are needed to determine whether LIN-23 directly targets WDR-23 and to define the molecular basis of this interaction. An equally important question concerns the context-dependent regulation of SKN-1 by LIN-23. Previous studies identified LIN-23 as a negative regulator of SKN-1 during embryogenesis, whereas our work demonstrates that it functions as a positive regulator during stress responses. Elucidating the mechanisms underlying this functional switch will provide valuable insights into the dynamic regulation of SKN-1 across distinct physiological contexts. Furthermore, given the large number of F-box proteins encoded in the *C. elegans* genome and the diverse roles of SCF ubiquitin ligase complexes in ubiquitin-mediated proteostasis, many aspects of LIN-23 regulation remain unknown. In particular, the upstream signals that control LIN-23 activation and substrate specificity during pathogen exposure warrant further investigation. Despite these outstanding questions, our study identifies LIN-23 as a novel positive regulator of SKN-1 during pathogen challenge and provides new insights into the mechanisms that enhance SKN-1-mediated protective responses, promoting host survival during infection.

## DATA AVAILABILITY

All data necessary for confirming the conclusions of this article are represented within the article. The raw data that underlies the figures is provided in Supplementary Table 3. The sources for all strains and reagents are provided in the Reagents Table.

## ACKNOWLEDGEMENTS

We thank the Caenorhabditis Genetics Center (CGC) for supplying strains, with support from the National Institutes of Health – Office of Research Infrastructure Programs (P40 OD010440). We thank Invivo Biosystems for aiding in the design and construction of our transgenic strains. We thank O. Karakuzu for intellectual contributions and A. Urrutia for technical support with this project.

## FIGURE LEGENDS

**Supplemental Fig. 1.**
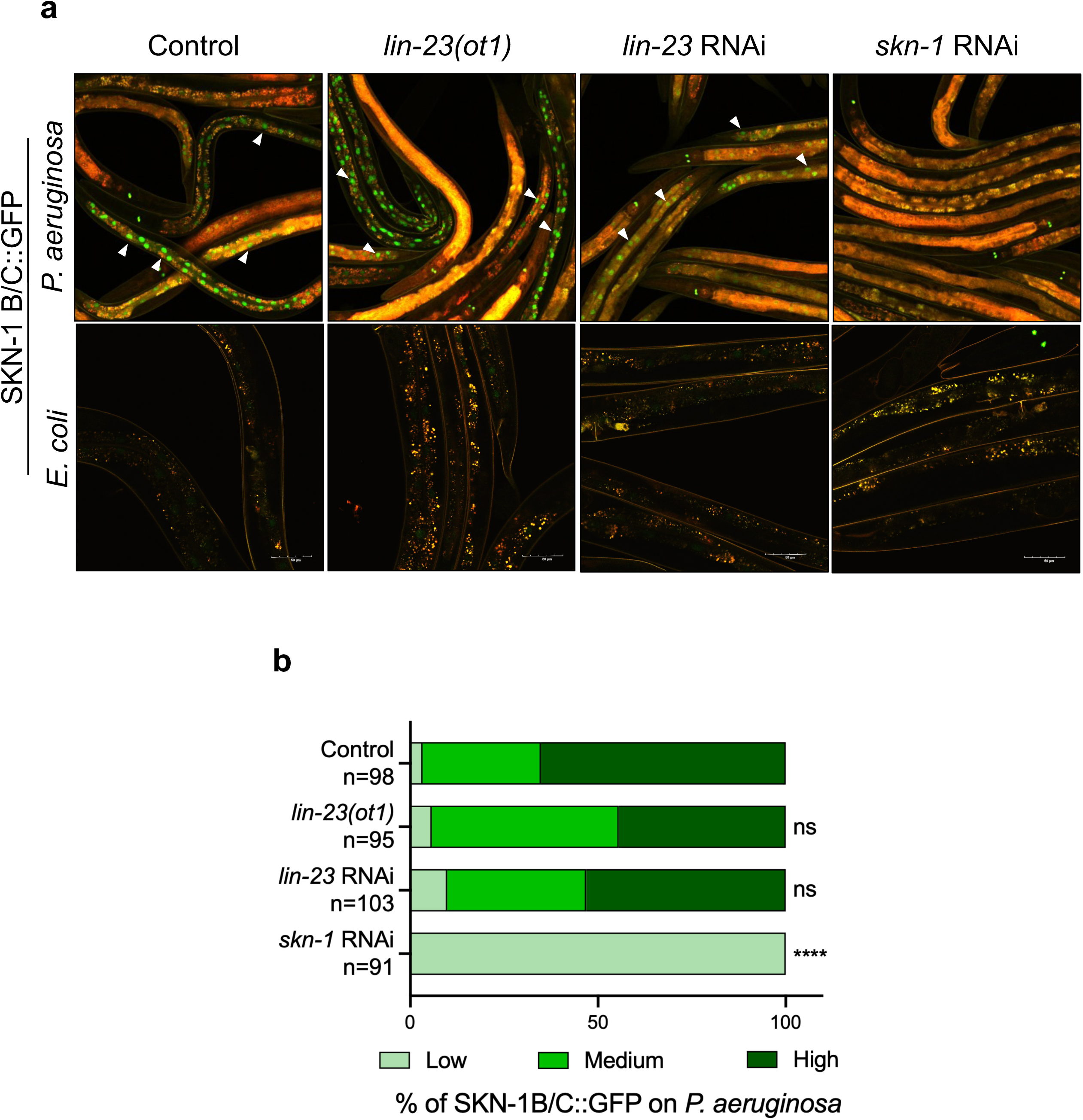
Effects of LIN-23 on SKN-1 nuclear localization during *P. aeruginosa* exposure. a) Wild-type and *lin-23(ot1)* worms containing an integrated SKN-1B/C::GFP transgene were exposed to *P. aeruginosa* or *E. coli*, and SKN-1B/C::GFP localization was examined by fluorescence microscopy. Representative intestinal nuclei exhibiting nuclear localization of SKN-1B/C::GFP are indicated by white arrows. Scale bars represent 50 μm. b) SKN-1C nuclear localization was scored based on the GFP signal and categorized as low, medium, or high. The percentage for each category was quantified, and the number of worms used in scoring each experimental condition is indicated (*n*). Levels of intestinal SKN-1B/C::GFP nuclear localization in *lin-23(ot1)* mutants and wild-type animals treated with *lin-23* or *skn-1* RNAi were compared to their respective controls. \*\*\*\**P* < 0.0001; ns, not significant.

**Supplemental Fig. 2.**
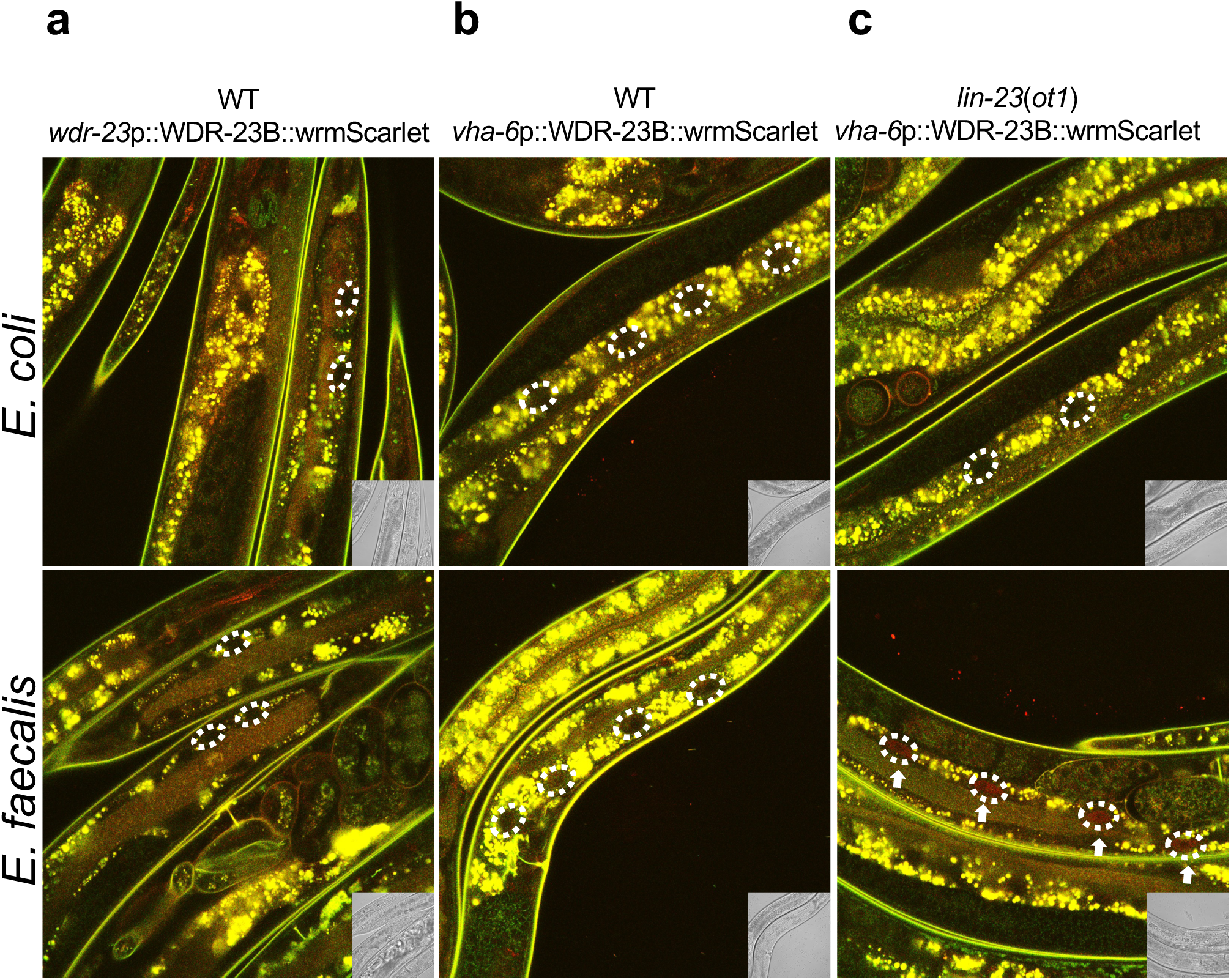
WDR-23B::wrmScarlet expressed under endogenous promoter. a) Representative images of WDR-23B::wrmScarlet expressed under the endogenous *wdr-23* promoter in wild-type background following exposure to *E. coli* or *E. faecalis*. Representative intestinal nuclei are outlined with white dashed lines. WDR-23B fluorescence was not detected under the endogenous promoter in either condition, likely due to low endogenous expression levels. b-c) Representative images of WDR-23B::wrmScarlet expressed under the intestine-specific *vha-6* promoter in wild-type and *lin-23(ot1)* backgrounds following exposure to *E. coli* or *E. faecalis*. <u>These images are duplicates of the data presented in the main text and are included here to facilitate side-by-side comparison between genotypes.</u>

## Supplemental Data: Western blot replicates

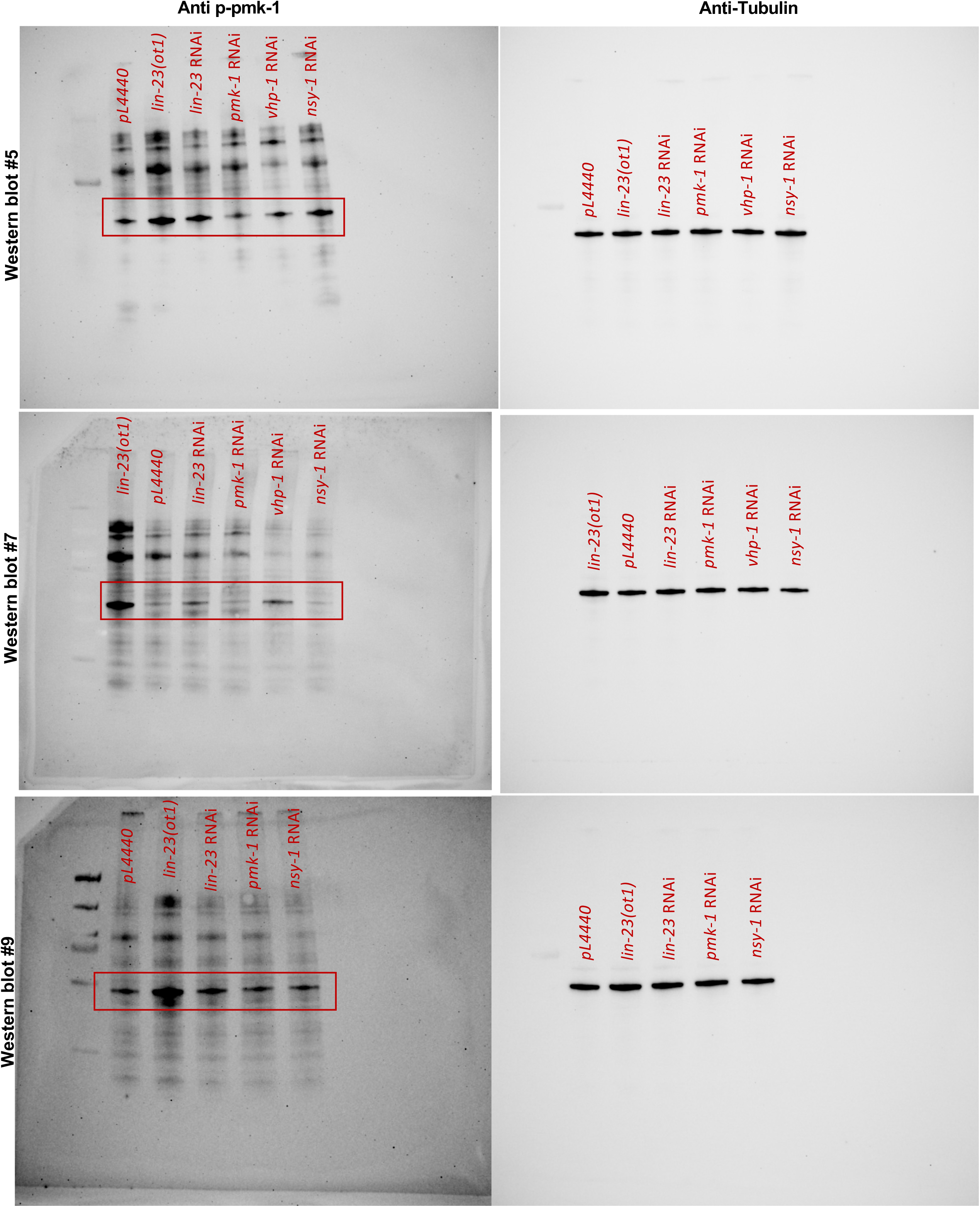

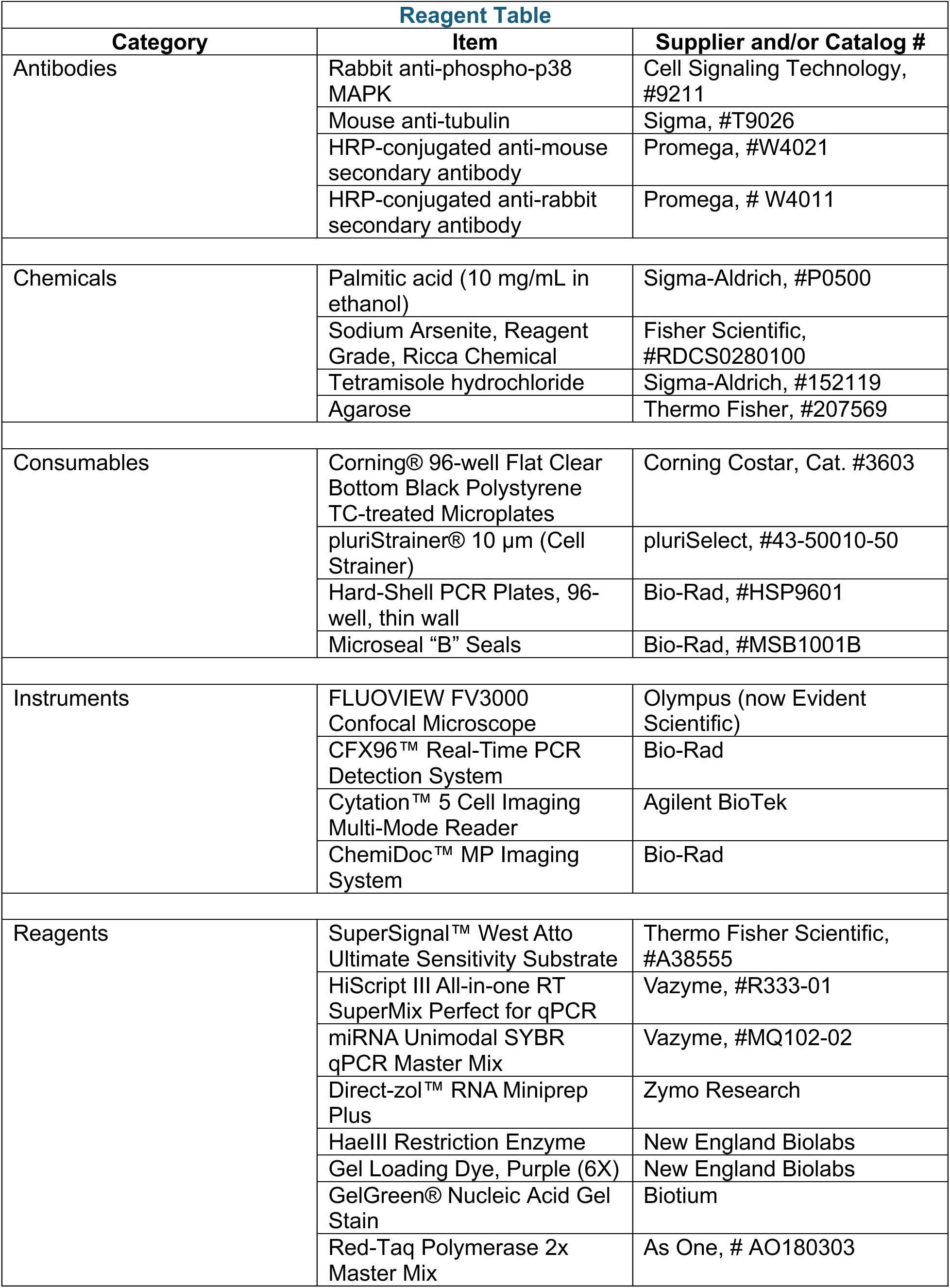
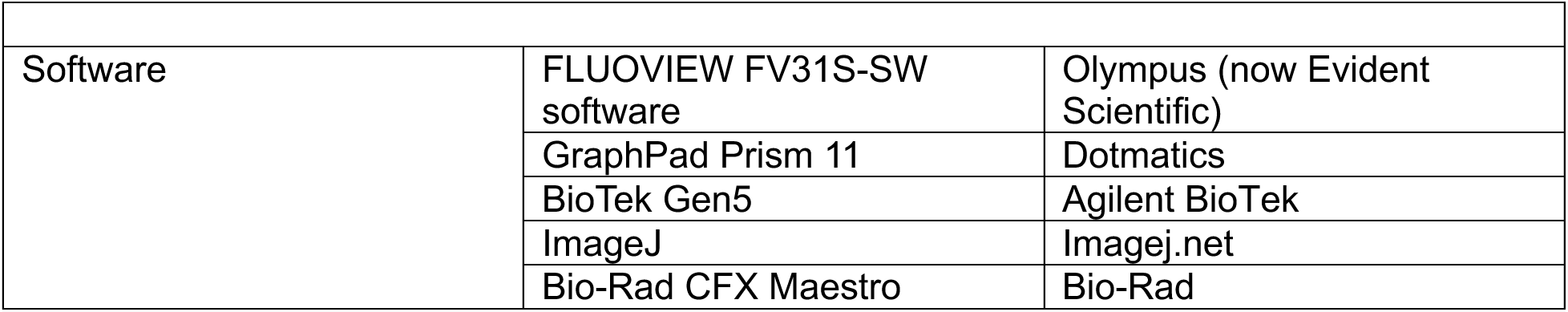

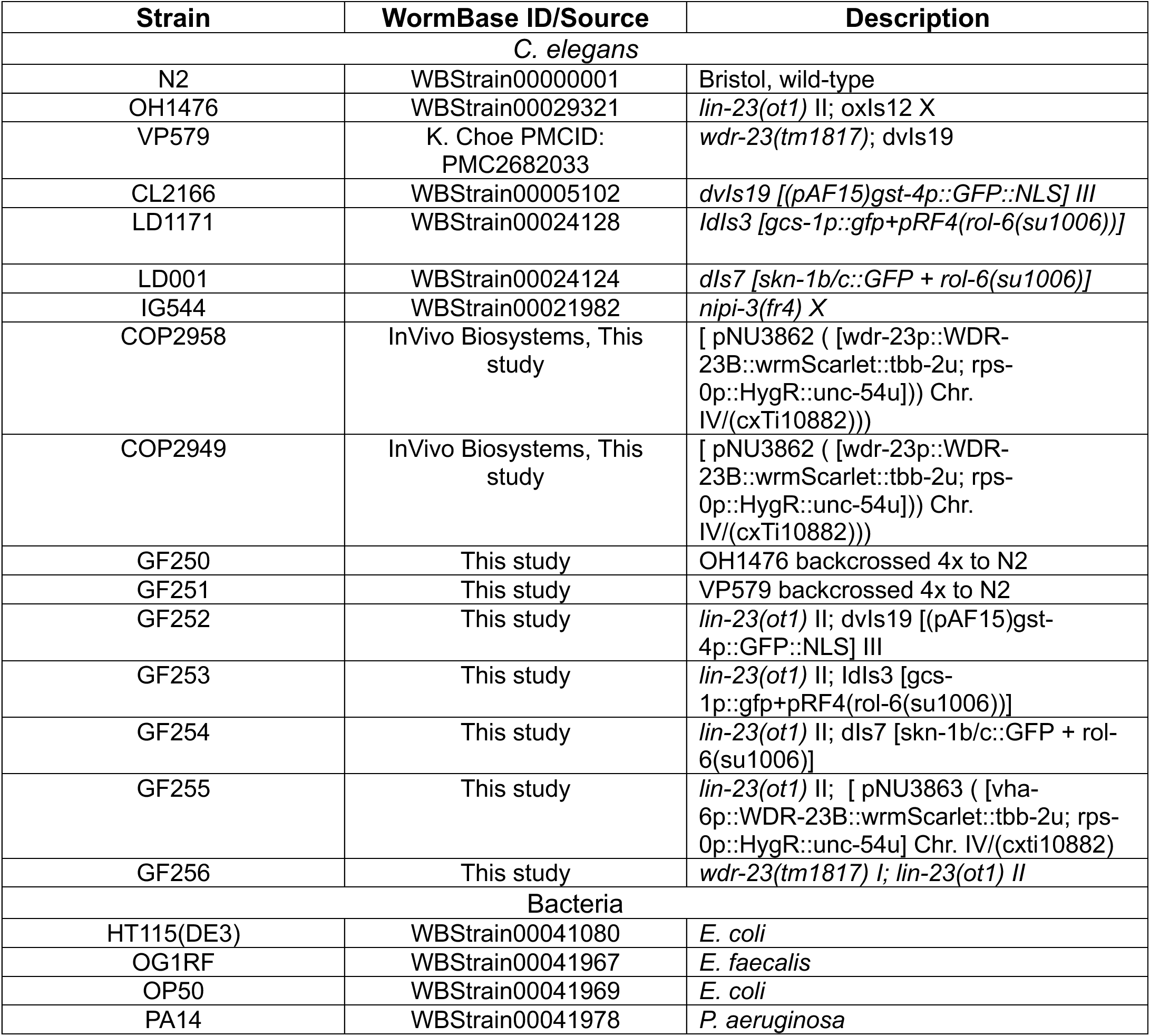
Supplementary Table 1.

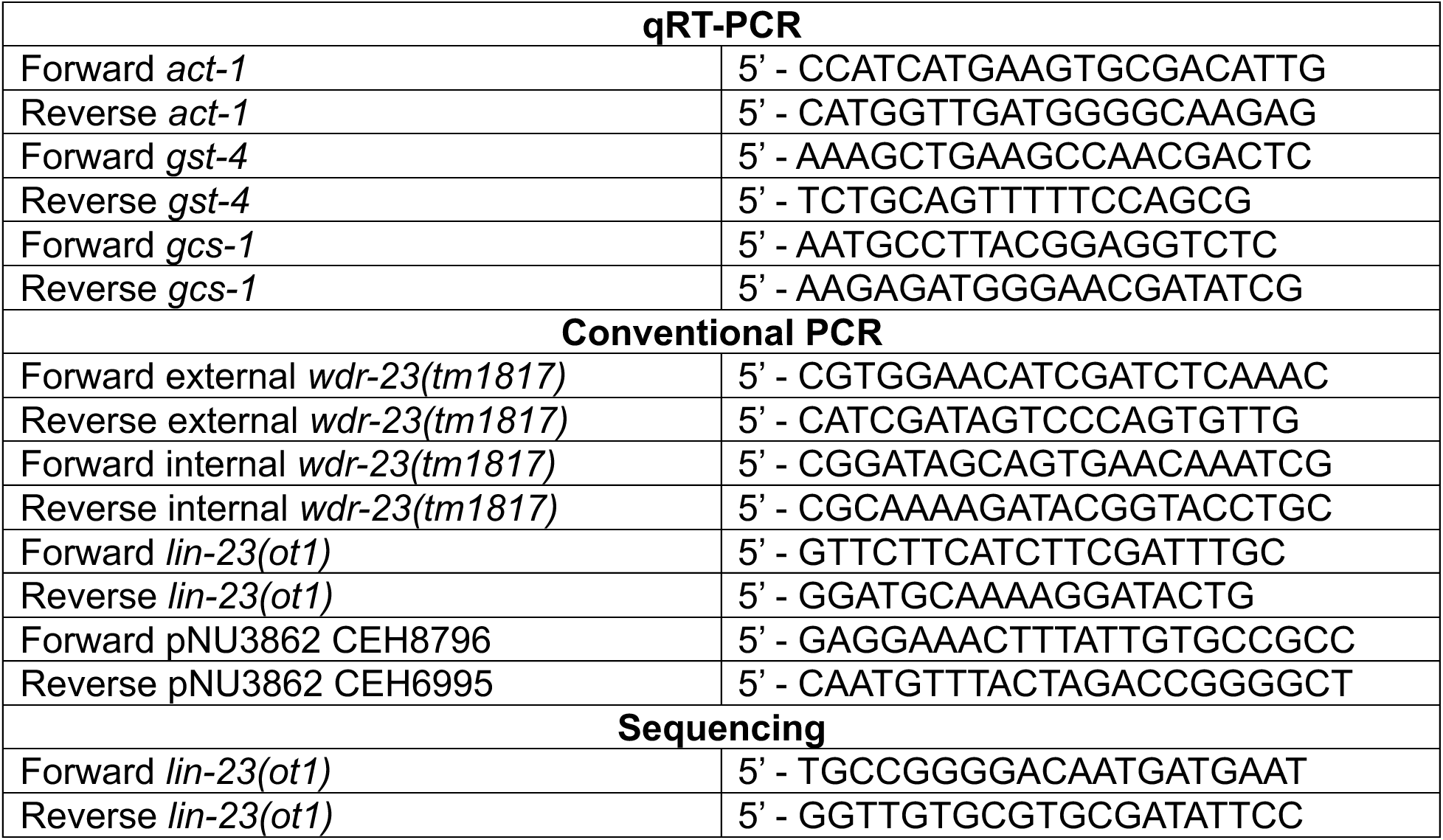
Supplementary Table 2.

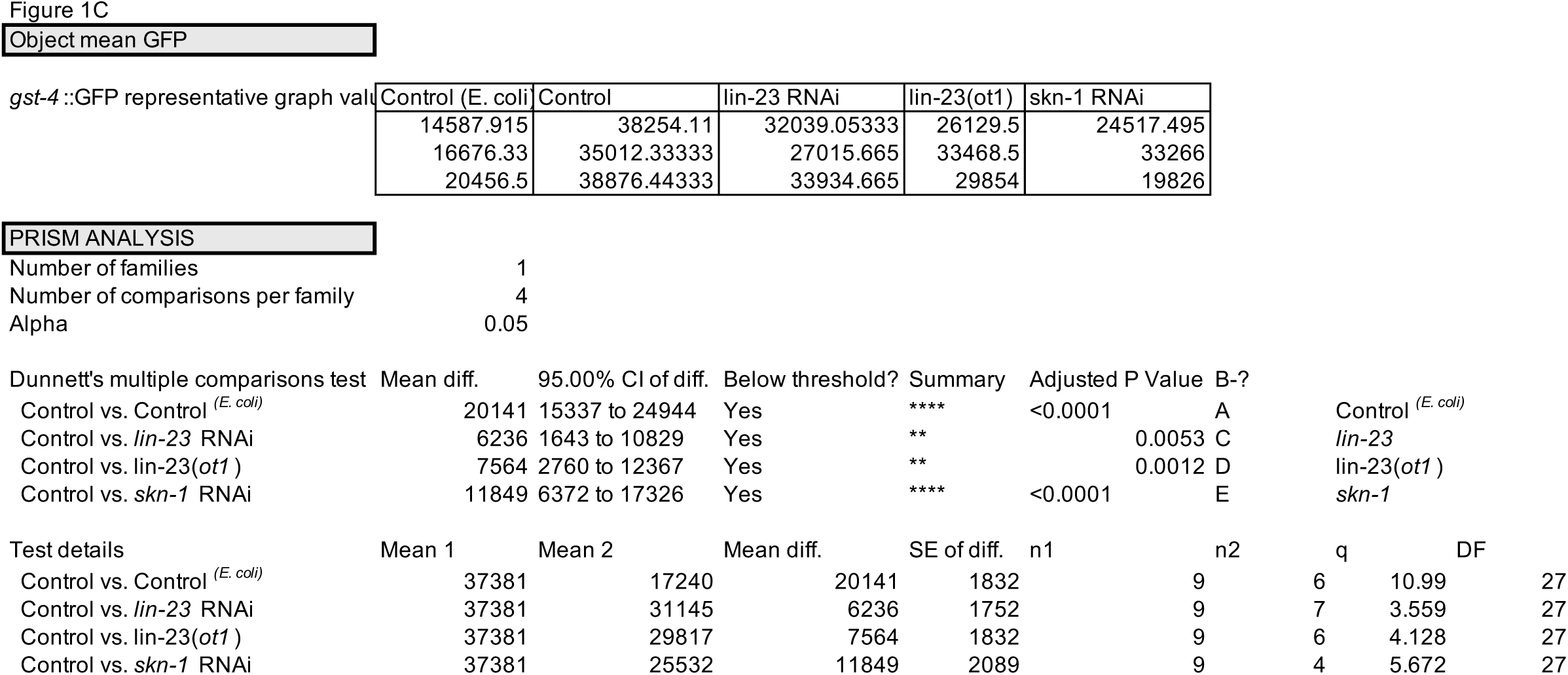

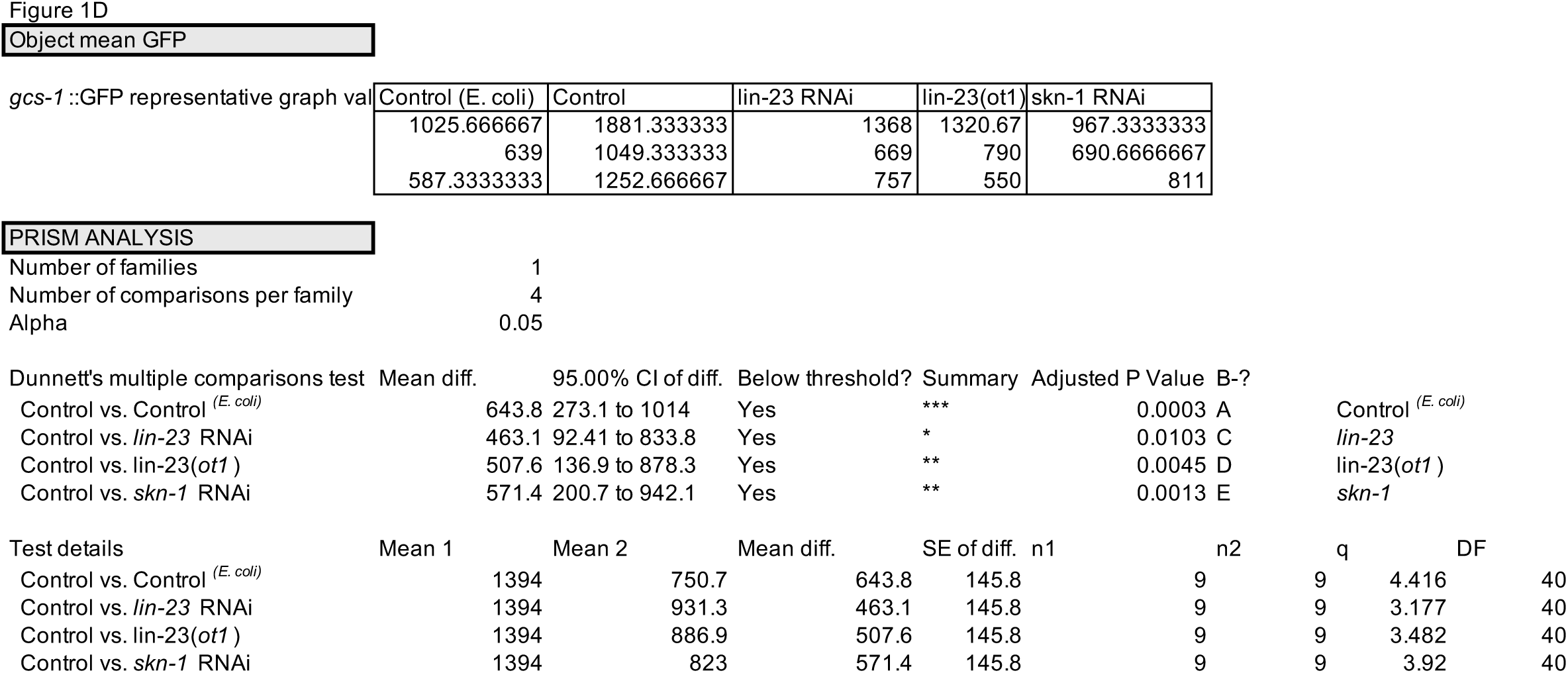

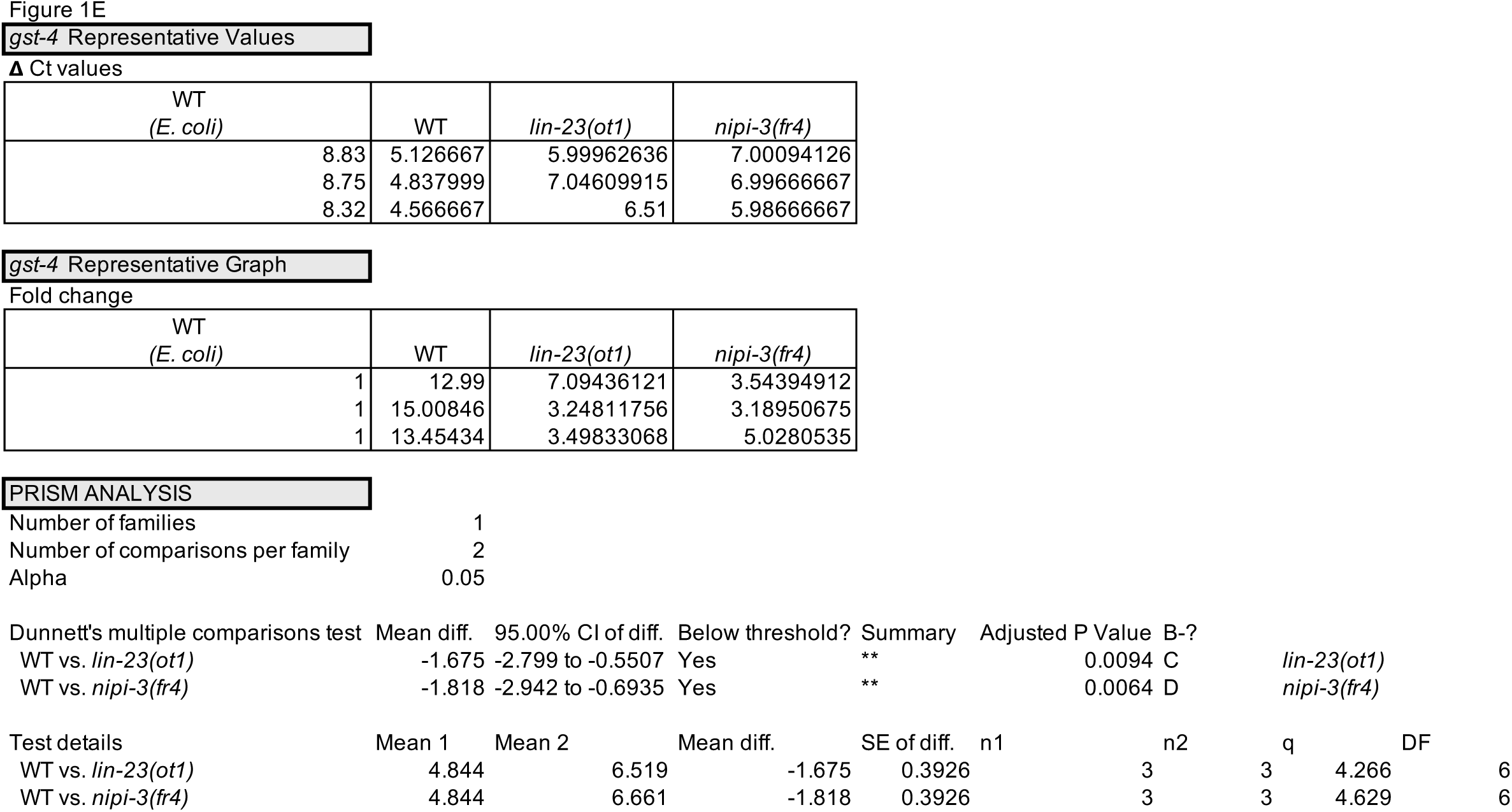

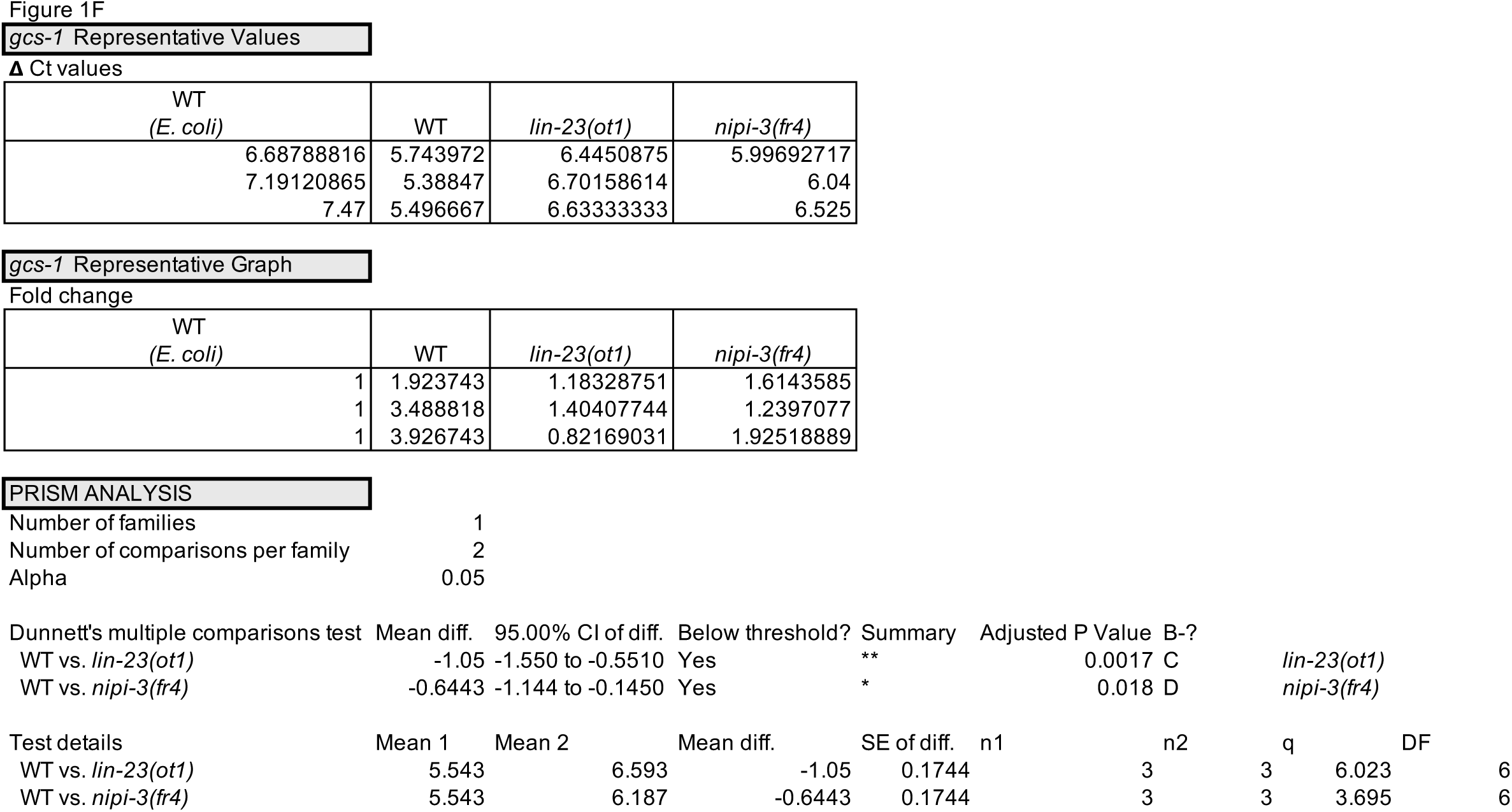

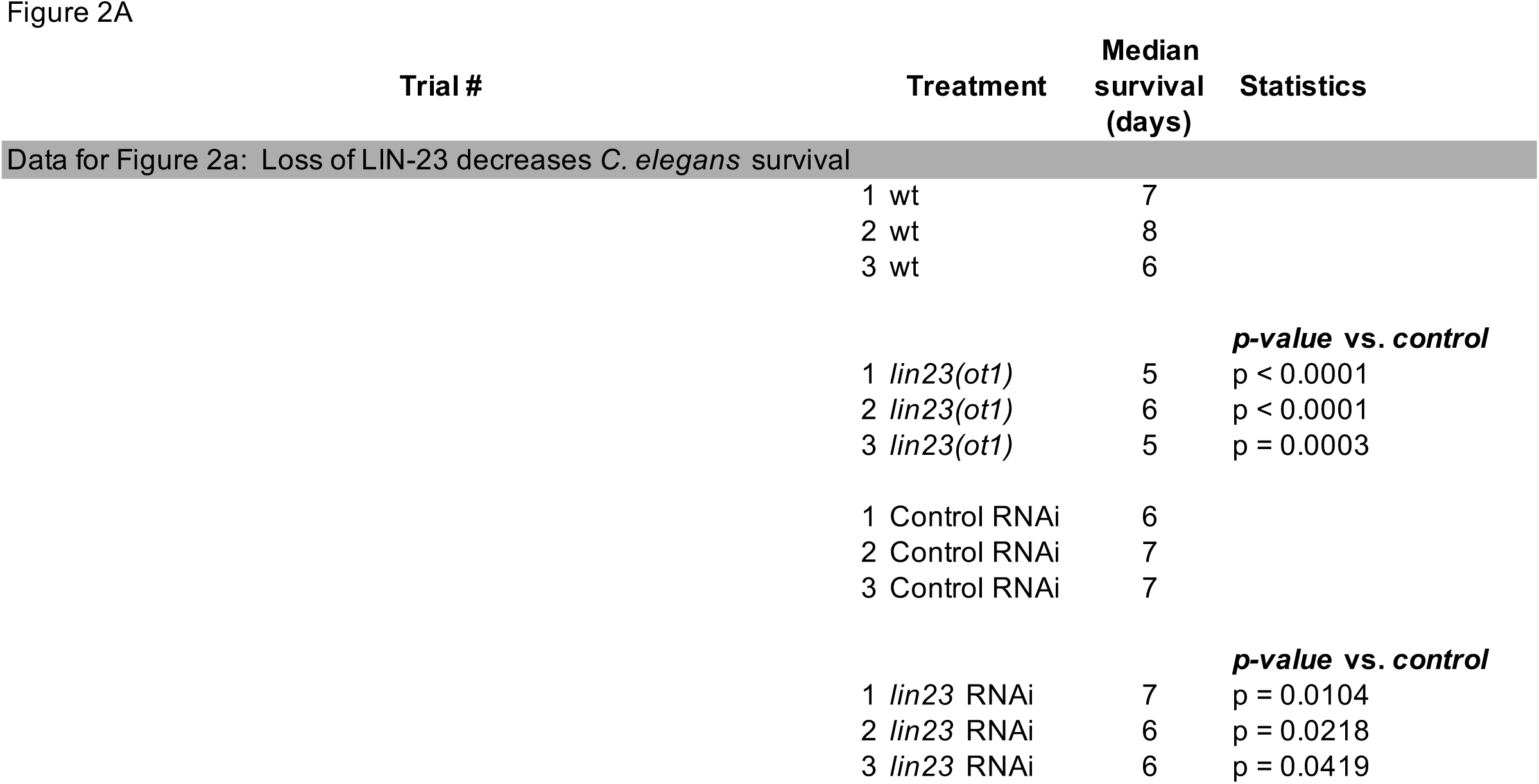

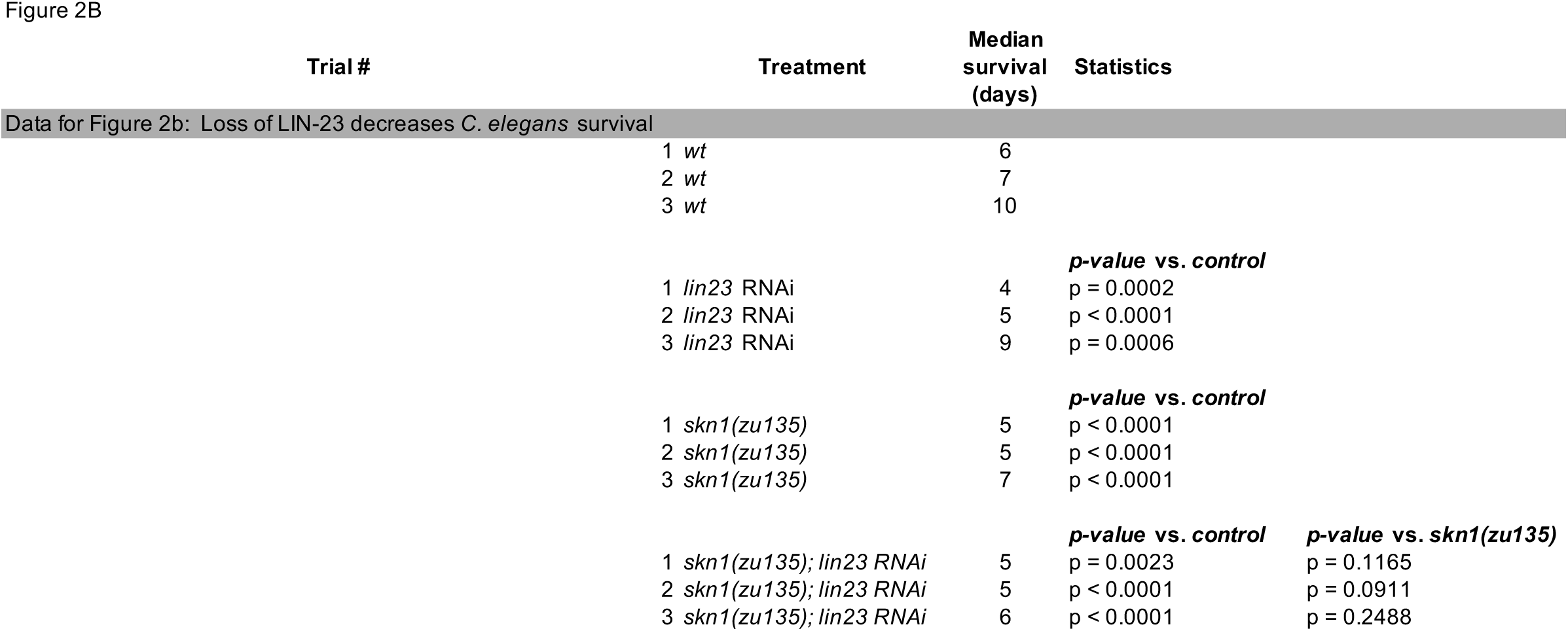

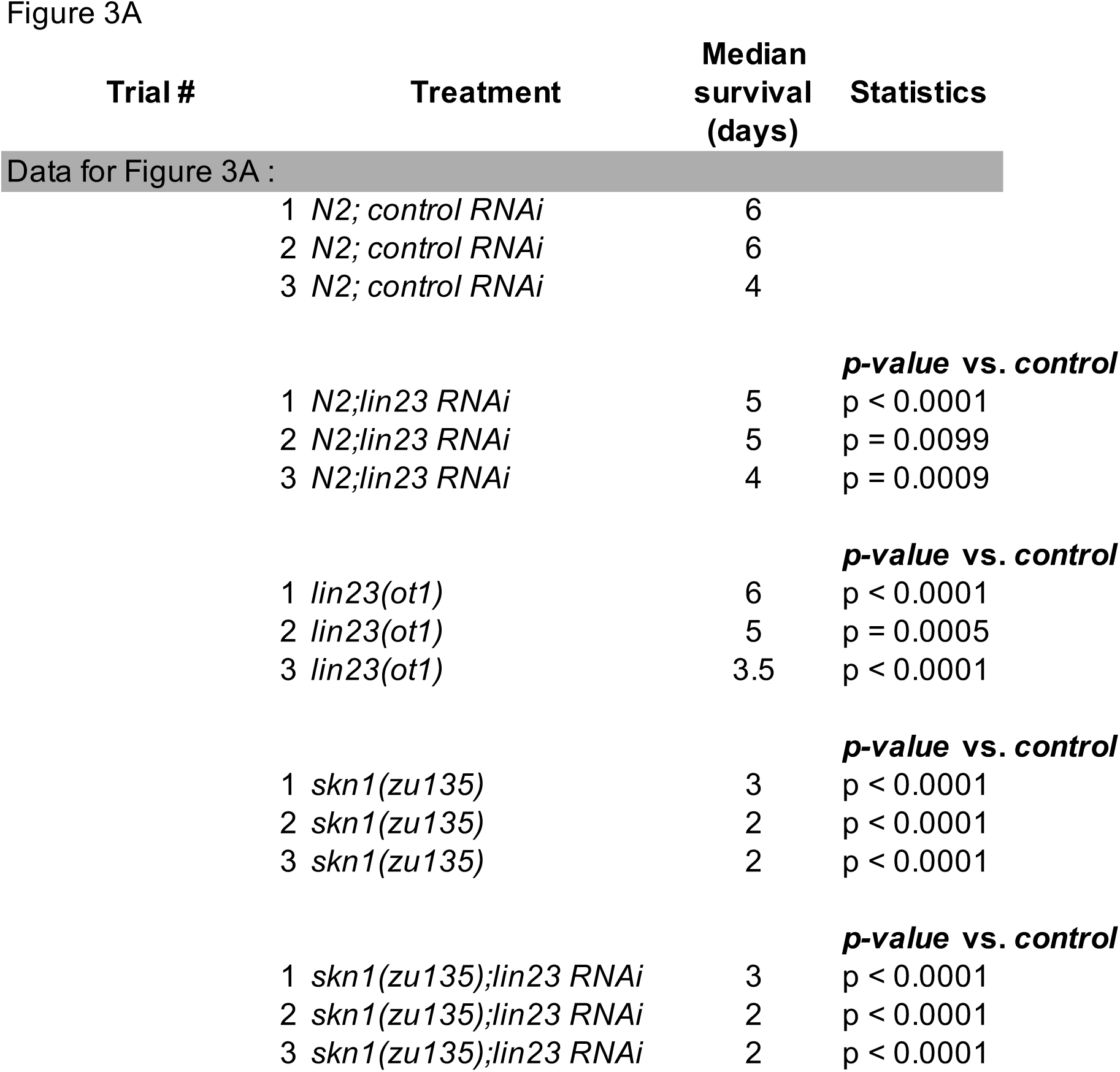

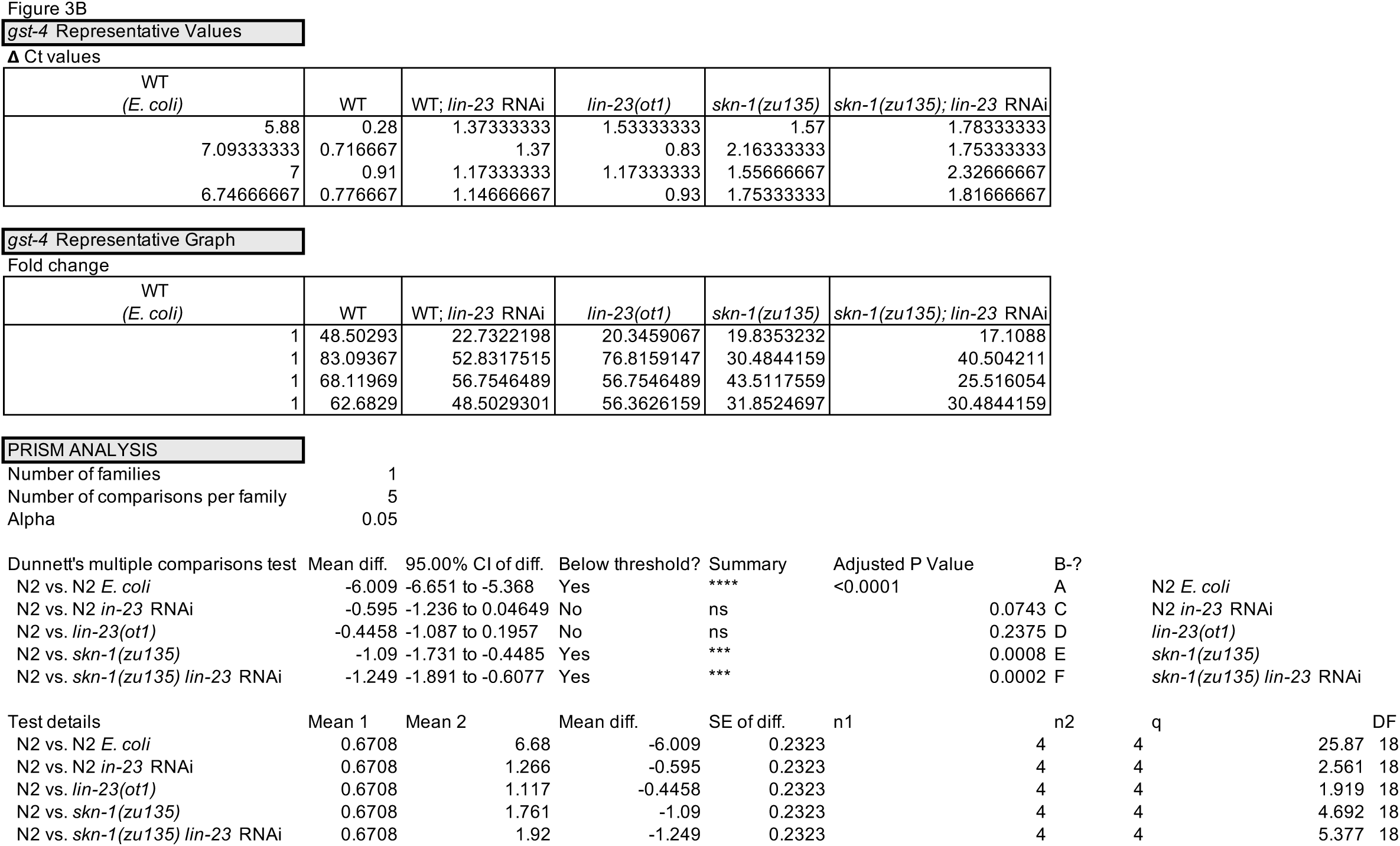

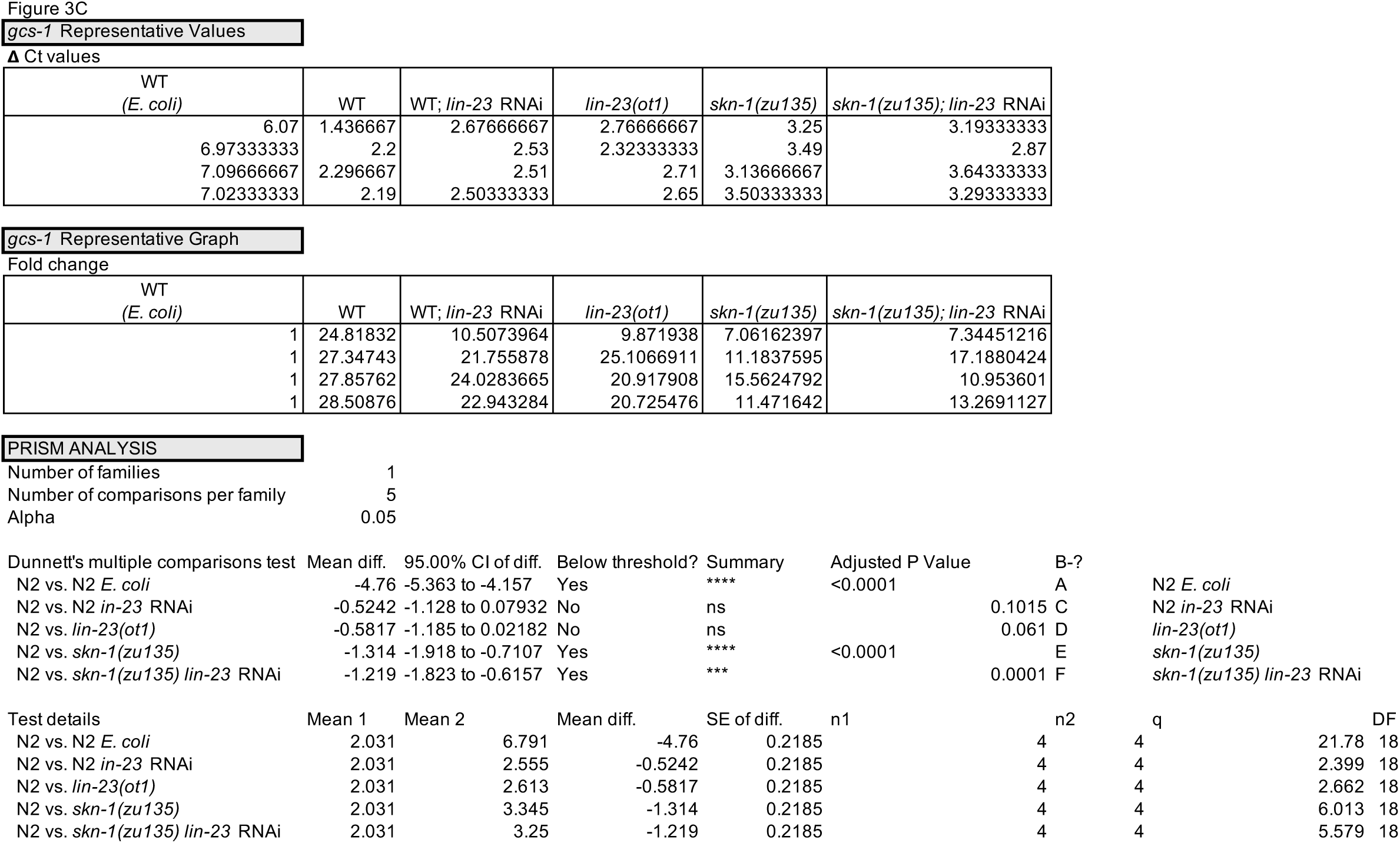

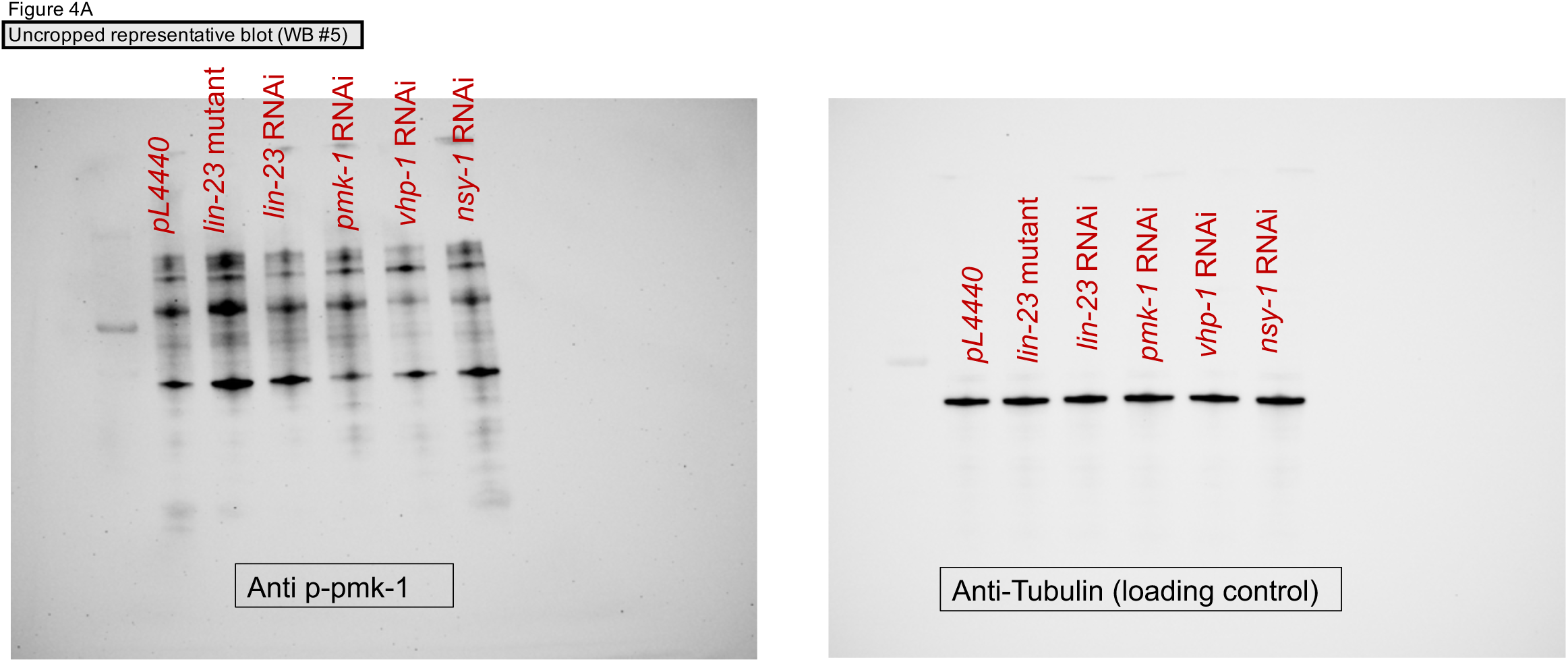

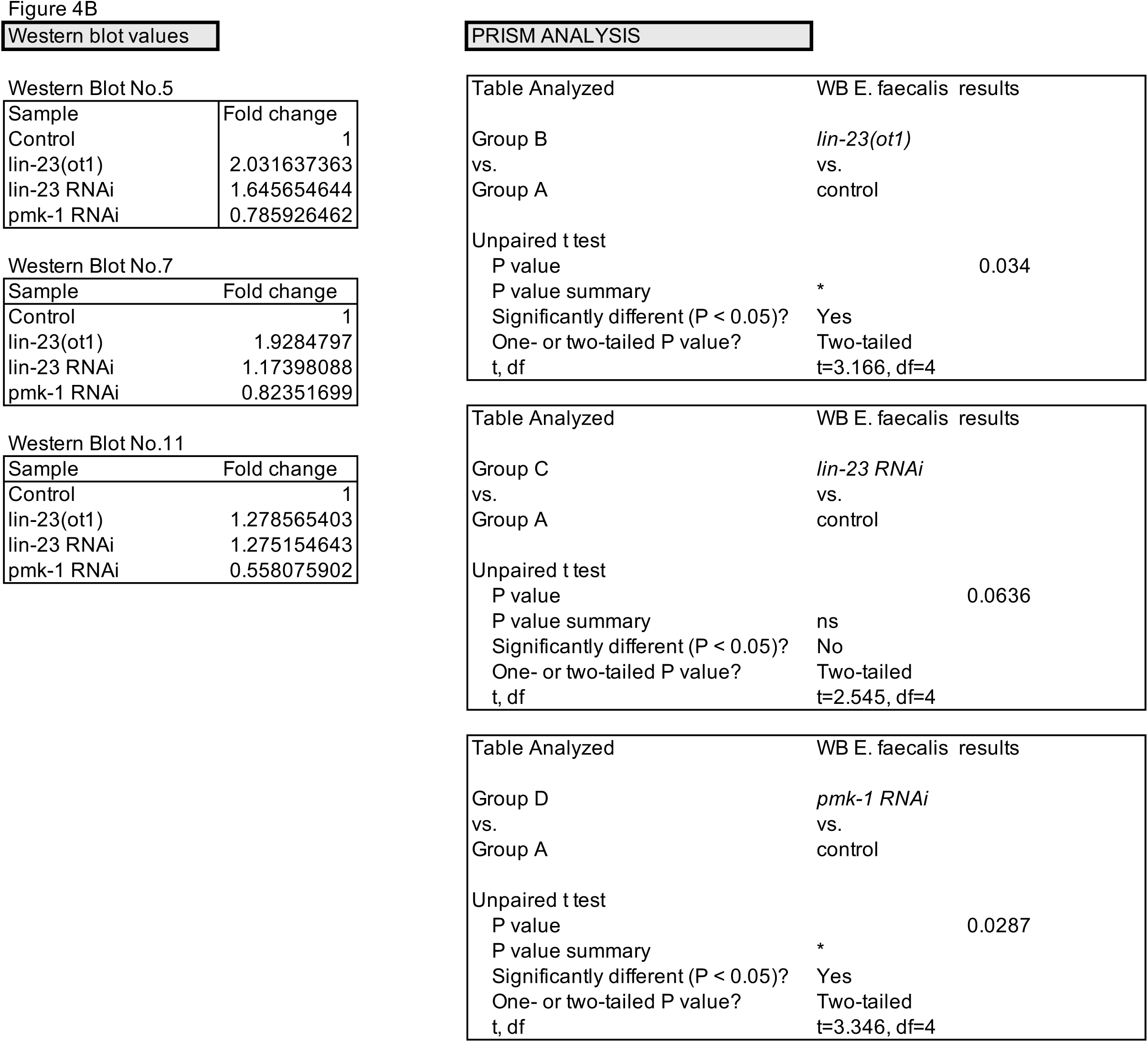

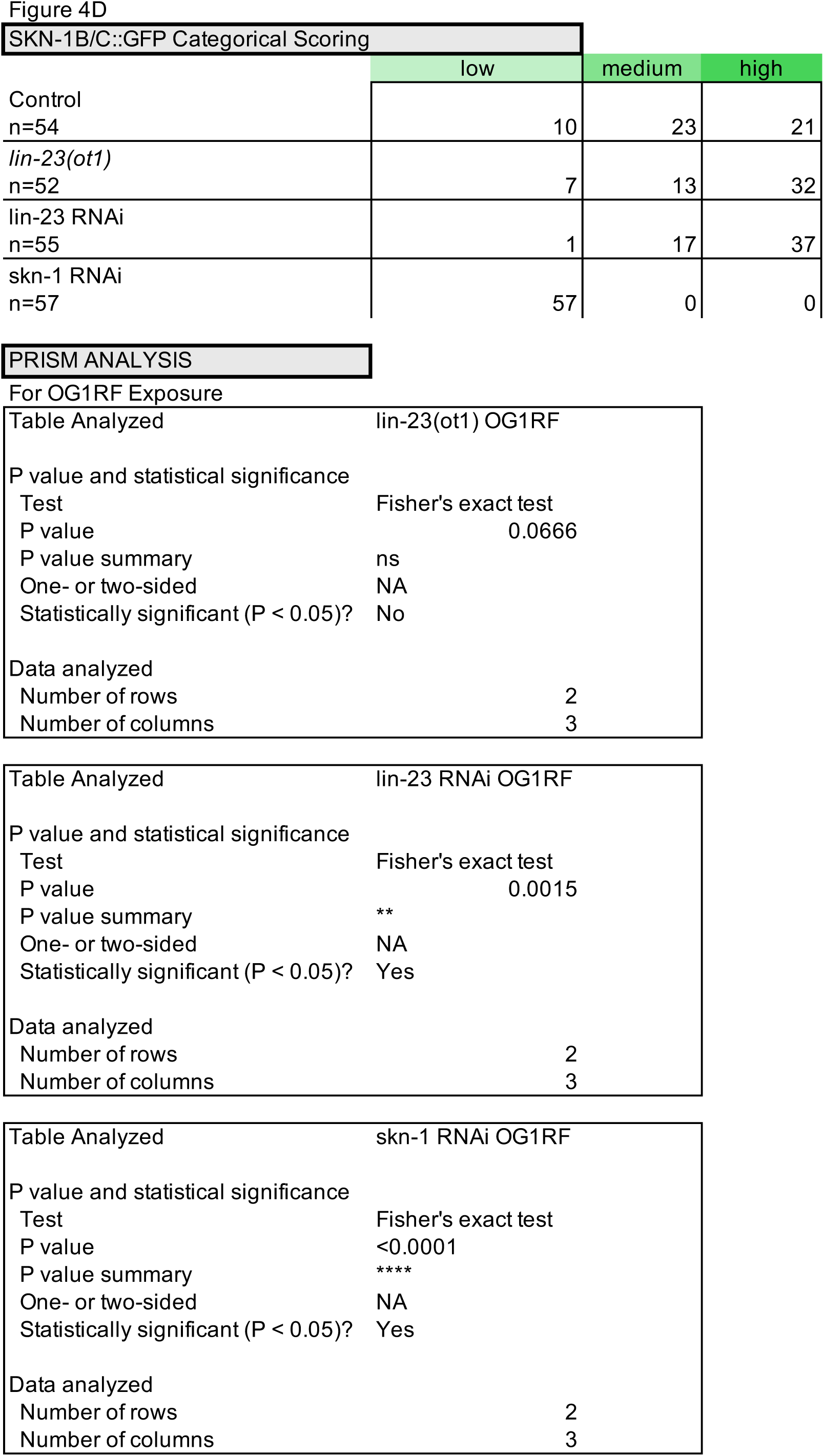

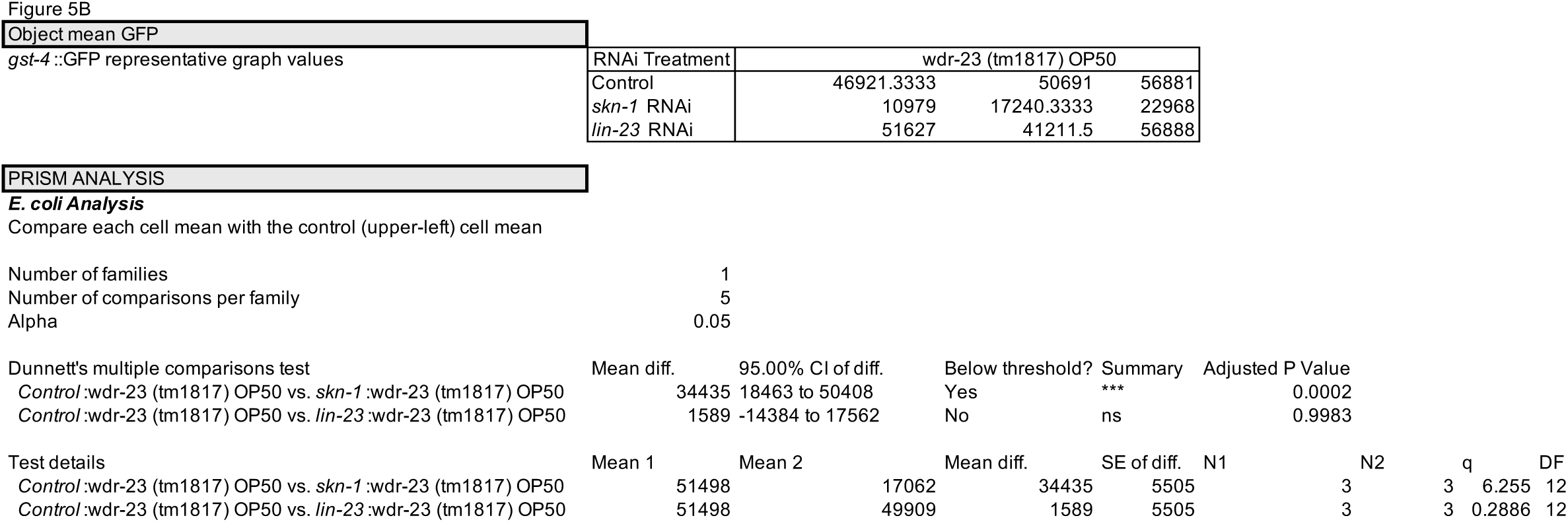

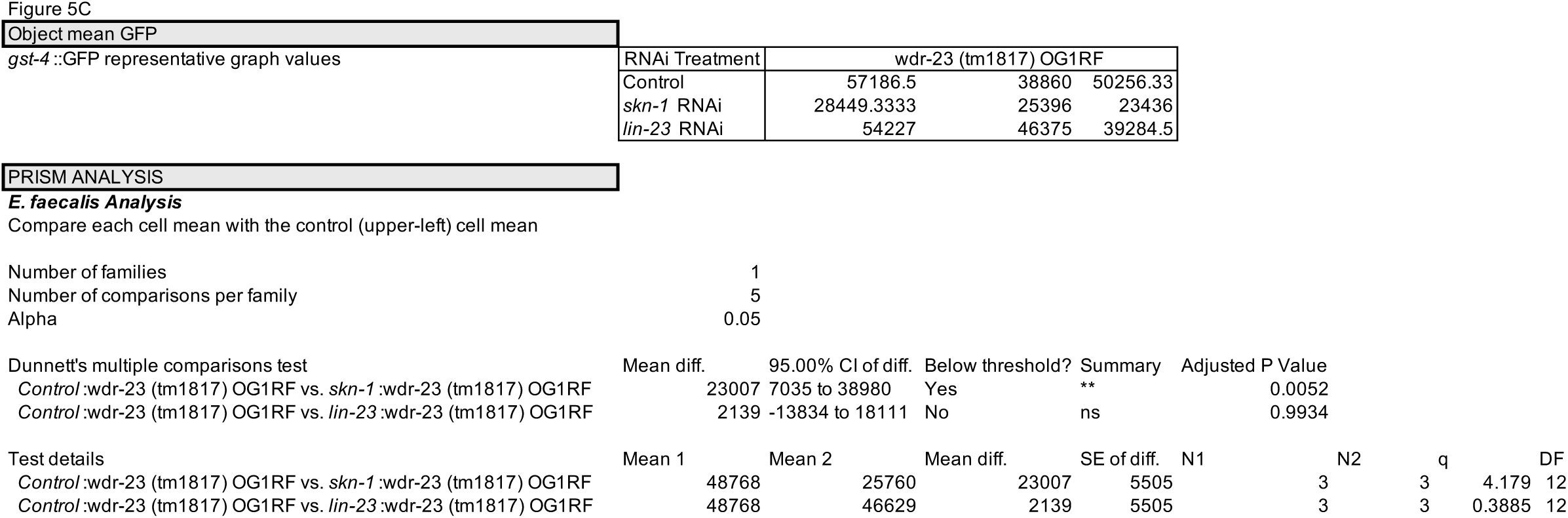

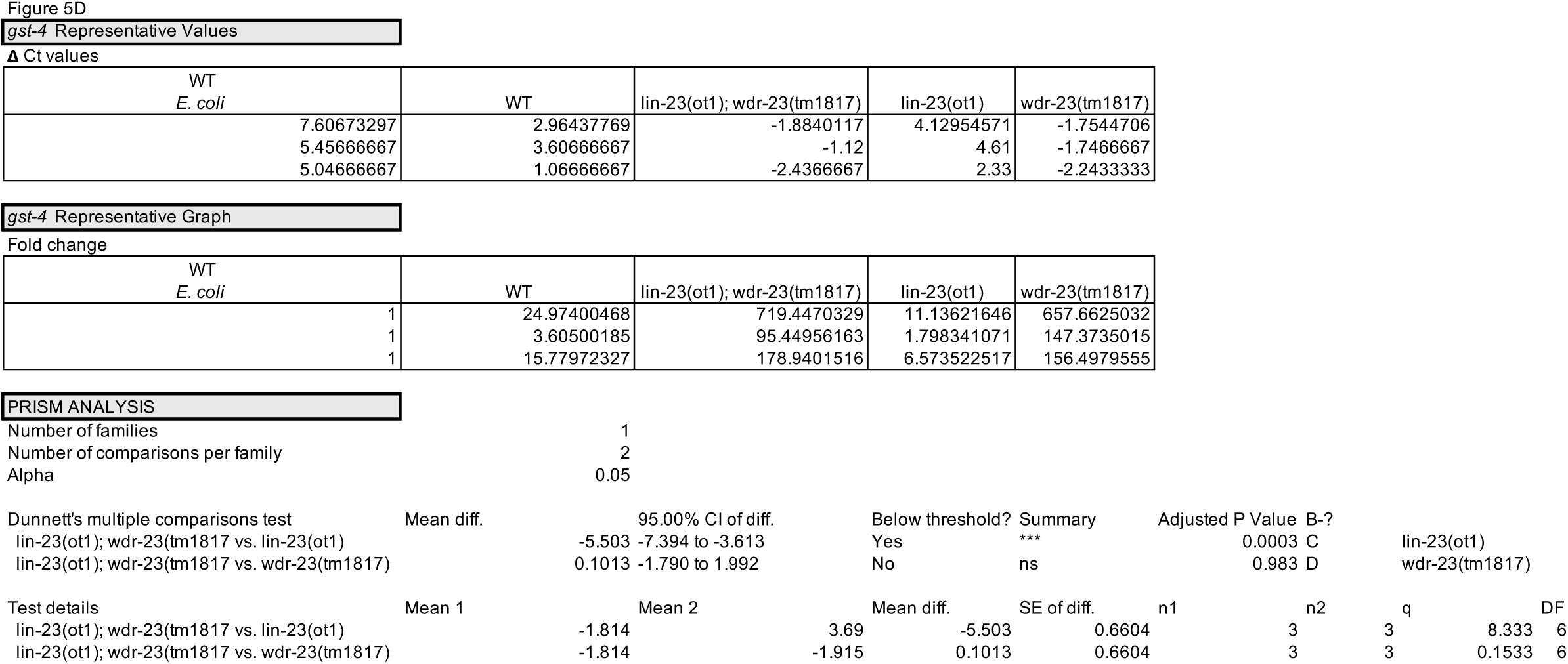

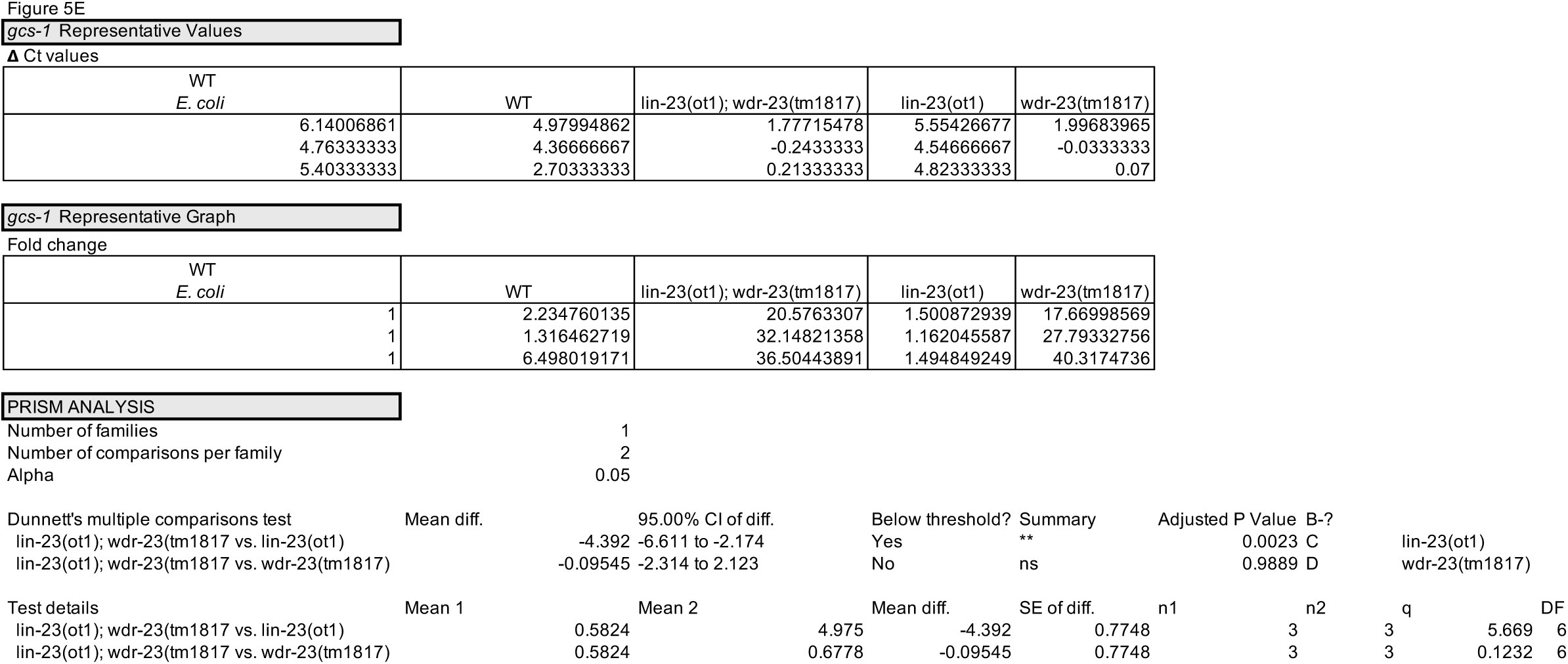

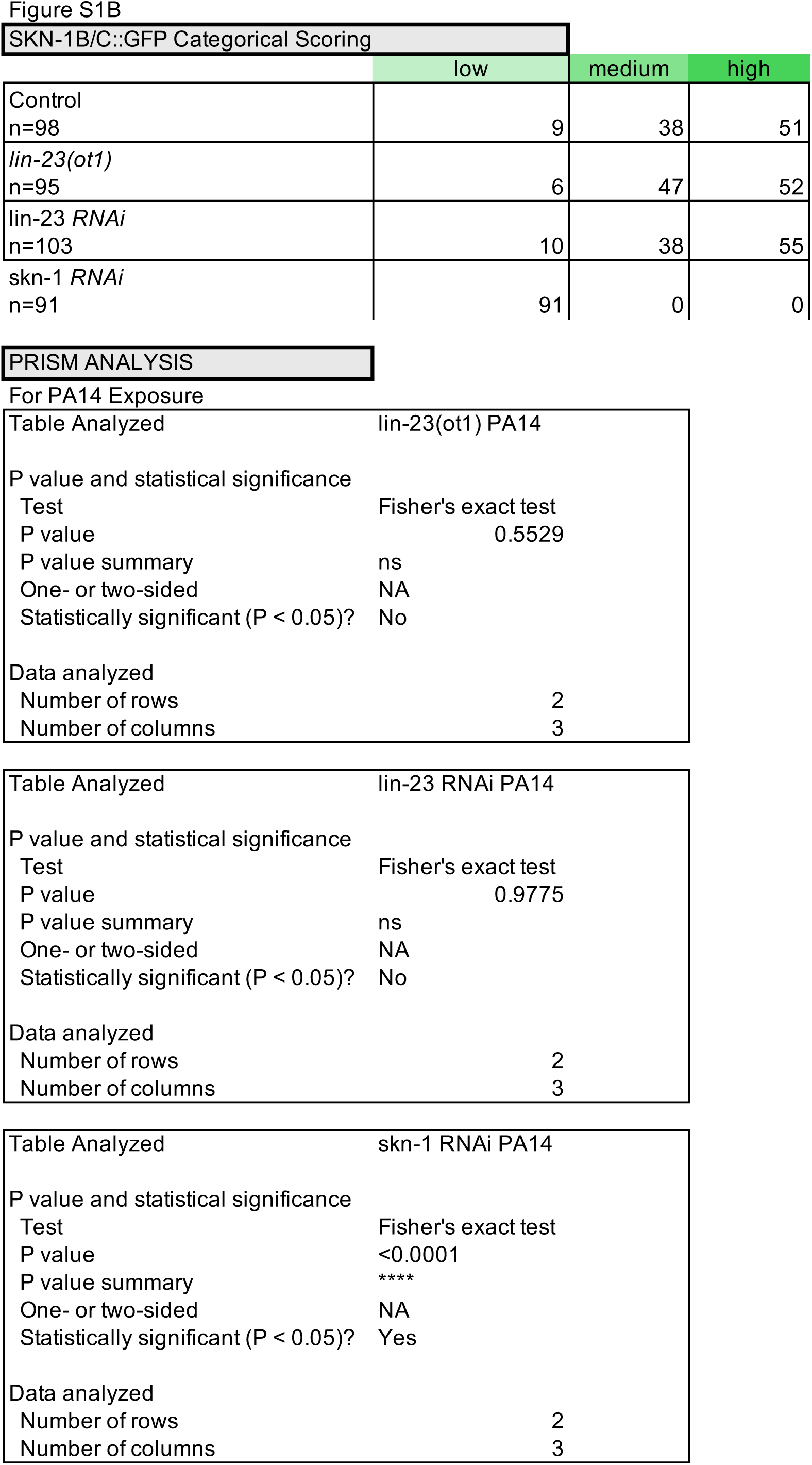

